# A Haplotype-resolved, Chromosome-scale Genome for *Malus domestica* Borkh. *‘*WA 38’

**DOI:** 10.1101/2024.01.10.574953

**Authors:** Huiting Zhang, Itsuhiro Ko, Abigail Eaker, Sabrina Haney, Ninh Khuu, Kara Ryan, Aaron B. Appleby, Brendan Hoffmann, Henry Landis, Kenneth Pierro, Noah Willsea, Heidi Hargarten, Alan Yocca, Alex Harkess, Loren Honaas, Stephen Ficklin

## Abstract

Genome sequencing for agriculturally important Rosaceous crops has made rapid progress both in completeness and annotation quality. Whole genome sequence and annotation gives breeders, researchers, and growers information about cultivar specific traits such as fruit quality, disease resistance, and informs strategies to enhance postharvest storage. Here we present a haplotype-phased, chromosomal level genome of *Malus domestica*, ‘WA 38’, a new apple cultivar released to market in 2017 as Cosmic Crisp ®. Using both short and long read sequencing data with a k-mer based approach, chromosomes originating from each parent were assembled and segregated. This is the ***first*** pome fruit genome fully phased into parental haplotypes in which chromosomes from each parent are identified and separated into their unique, respective haplomes. The two haplome assemblies, ‘Honeycrisp’ originated HapA and ‘Enterprise’ originated HapB, are about 650 Megabases each, and both have a BUSCO score of 98.7% complete. A total of 53,028 and 54,235 genes were annotated from HapA and HapB, respectively. Additionally, we provide genome-scale comparisons to ‘Gala’, ‘Honeycrisp’, and other relevant cultivars highlighting major differences in genome structure and gene family circumscription. This assembly and annotation was done in collaboration with the American Campus Tree Genomes project that includes ‘WA 38’ (Washington State University), ‘d’Anjou’ pear (Auburn University), and many more. To ensure transparency, reproducibility, and applicability for any genome project, our genome assembly and annotation workflow is recorded in detail and shared under a public GitLab repository. All software is containerized, offering a simple implementation of the workflow.

## Introduction

For economically important crop species, having full-resolution reference genomes aids in the understanding of traits associated with commodity quality, disease resistance, long-term storage, and shelf life. Apple (*Malus domestica*) is the number one consumed fruit in the United States, with a Farm-Gate Revenue of $3.2 billion in the U.S. (USApple, 2024), and $78 billion globally (FGN, 2020). There are over 7,000 apple varieties grown world wide (Washington Apple Commission, 2024), each with unique colors, flavors, and textures (N.C. Cooperative Extension, n.d.). Therefore, a single genome is unlikely to capture the complexity of all cultivars within this highly heterozygous species (Li *et al*. 2022b; Zhang *et al*. 2022). One such cultivar is ‘WA 38’, commercially released as Cosmic Crisp® in 2017 by the Pome Fruit Breeding Program at Washington State University’s (WSU) Tree Fruit Research and Extension Center (Figure 1 A, B) and has reached the top 10 best selling apple cultivars in the United States (Truscott, 2023). ‘WA 38’ is a cross between ‘Honeycrisp’ and ‘Enterprise’, made using classical breeding methods in 1997. One parent, ‘Honeycrisp’, is well-known for its crisp texture, firmness retention in storage, disease resistance, and cold hardiness, but is highly susceptible to production and postharvest issues (Khan *et al*. 2022). The other parent, ‘Enterprise’, is an easy-to-grow cultivar that has extended postharvest storage capabilities, however it is not widely cultivated commercially due to its less desirable eating quality (Crosby *et al*. 1994). Their resulting cross has been met with favorable reviews for its appealing color, texture, flavor, cold hardiness, long-term storage capabilities (>1 year), and scab resistance (Evans *et al*. 2012). However, it inherited undesirable traits as well, such as a propensity for physiological symptoms that may be related to mineral imbalances (Sallato *et al*. 2021; Sheick *et al*. 2023), maturity at harvest (Serra et al 2023), and an ‘off’ flavor that has been brought up by consumers that may be the result of improper picking times, crop load management, handling/packing practices or other post harvest processes (Mendoza, M., Hanrahan, I., & Bolaños, G., 2020). Most concerning is green spot (Figure 1 C, D, and F), a corking disorder that seems to be unique to ‘WA 38’, but with etiology similar to disorders associated with mineral imbalances such as bitter pit and drought spot (Sheick *et al*. 2022, 2023). The propensity for and cause of physiological disorders often differs on a cultivar-by-cultivar basis (Pareek 2019), and a genetic basis for such predispositions is likely (Liebhard *et al*. 2003; Johnston and Brookfield 2012; Di Guardo *et al*. 2013; Lum *et al*. 2016). Thus, improved resolution of cultivar-specific genomic differences is critical for advancing our understanding of how economically important traits, such as physiological disorders, are inherited and how they can be managed more efficiently.

**Figure 1.**
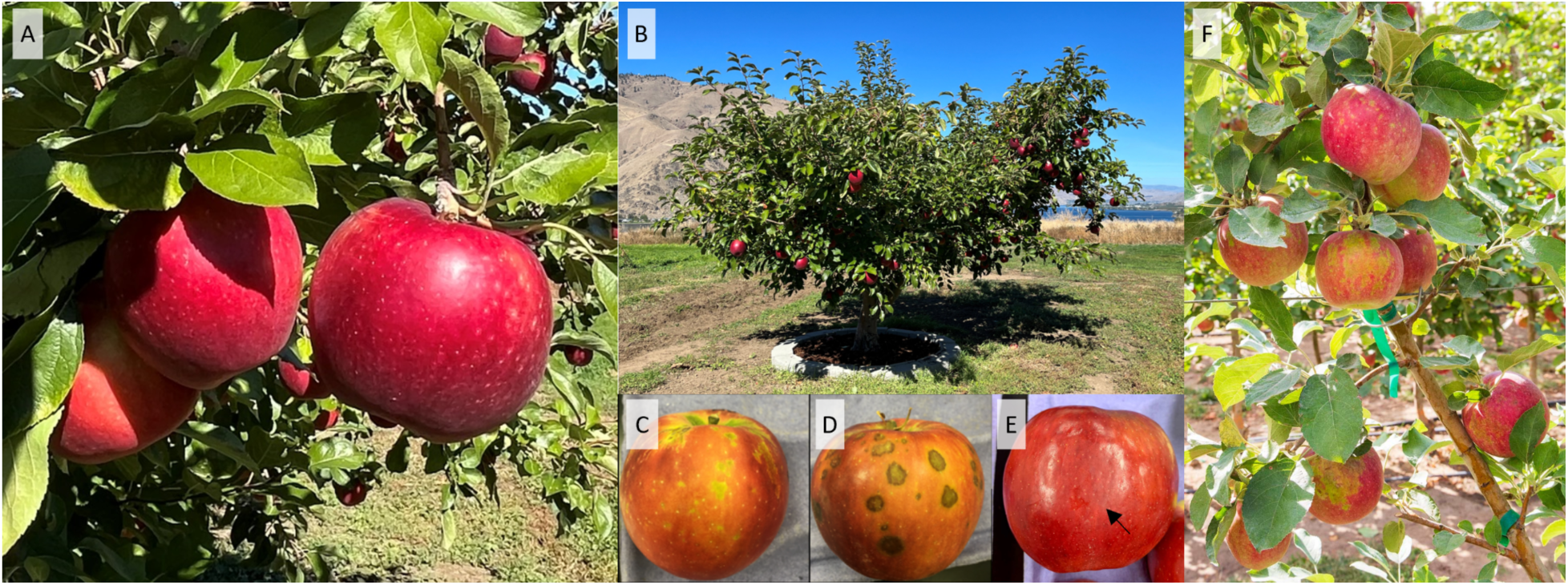
‘WA 38’, a cultivar of apple developed by the Washington State University Apple Breeding Program (a cross between ‘Honeycrisp’ and ‘Enterprise’), marketed as Cosmic Crisp®. **A)** ‘WA 38’ apples ready for harvest on the mother tree, located at the WSU and USDA-ARS Columbia View Research Orchard near Orondo, WA, USA. **B**) The ‘WA 38’ mother tree. **C** & **D**) Green spot, a corking disorder which results in green blemishes on the fruit peel and brown, corky cortex tissue. Symptom severity generally increases during fruit maturation and time in storage, resulting in cullage. **E**) Natural peel greasiness as a result of more advanced maturity at harvest can interfere with artificial waxes applied in the packinghouse after removal from postharvest storage, creating unappealing, dull spots. **F**) Green Spot symptoms can begin to appear while fruit is still developing on the tree. Photo Credits: A&B: Heidi Hargarten/USDA-ARS; C&D: Bernardita Sallato/WSU; E: Carolina Torres/WSU; F: Ross Courtney/Good Fruit Grower.

To develop full resolution reference genomes of superior quality, having skilled bioinformaticians is required. To train the next generation of bioinformaticians for agricultural genomic research, a national effort spearheaded by Auburn University, HudsonAlpha Institute for Biotechnology, and Washington State University was started in 2021 - The American Campus Tree Genomes project (ACTG). ACTG aims to break through institutional barriers that have traditionally prevented many students from accessing valuable, hands-on research projects and experience in bioinformatics (Sharman, S, n.d.). To accomplish this goal, a course has been developed to involve students in genome projects from inception, through analysis, to publication (Harkess, 2022). During the course, students learn genome assembly and annotation workflows using the raw sequence data from genomes of beloved trees (e.g., Toomer’s oak and ‘d’Anjou’ pear (Yocca *et al*. 2024) at Auburn University, Sabal palm at University of South Carolina) and are listed as authors on the final publication. The ‘WA 38’ genome introduced here was developed through ACTG by students from Washington State University, presenting three major outcomes: 1) a fully annotated, chromosomal level, haplotype-resolved genome of ‘WA 38’ utilizing PacBio HiFi, Dovetail Omni-C, and Illumina DNA and RNA sequencing data, 2) identification of unique regions of interest using a comparative genomics approach with other economically important *M. domestica* cultivars including ‘Gala’, ‘Fuji’, and ‘Honeycrisp’, and 3) establishment of a containerized, reproducible, flexible, high performance computing workflow for complete genome assembly and annotation (Figure 2, Supplemental Figure 1).

**Figure 2.**
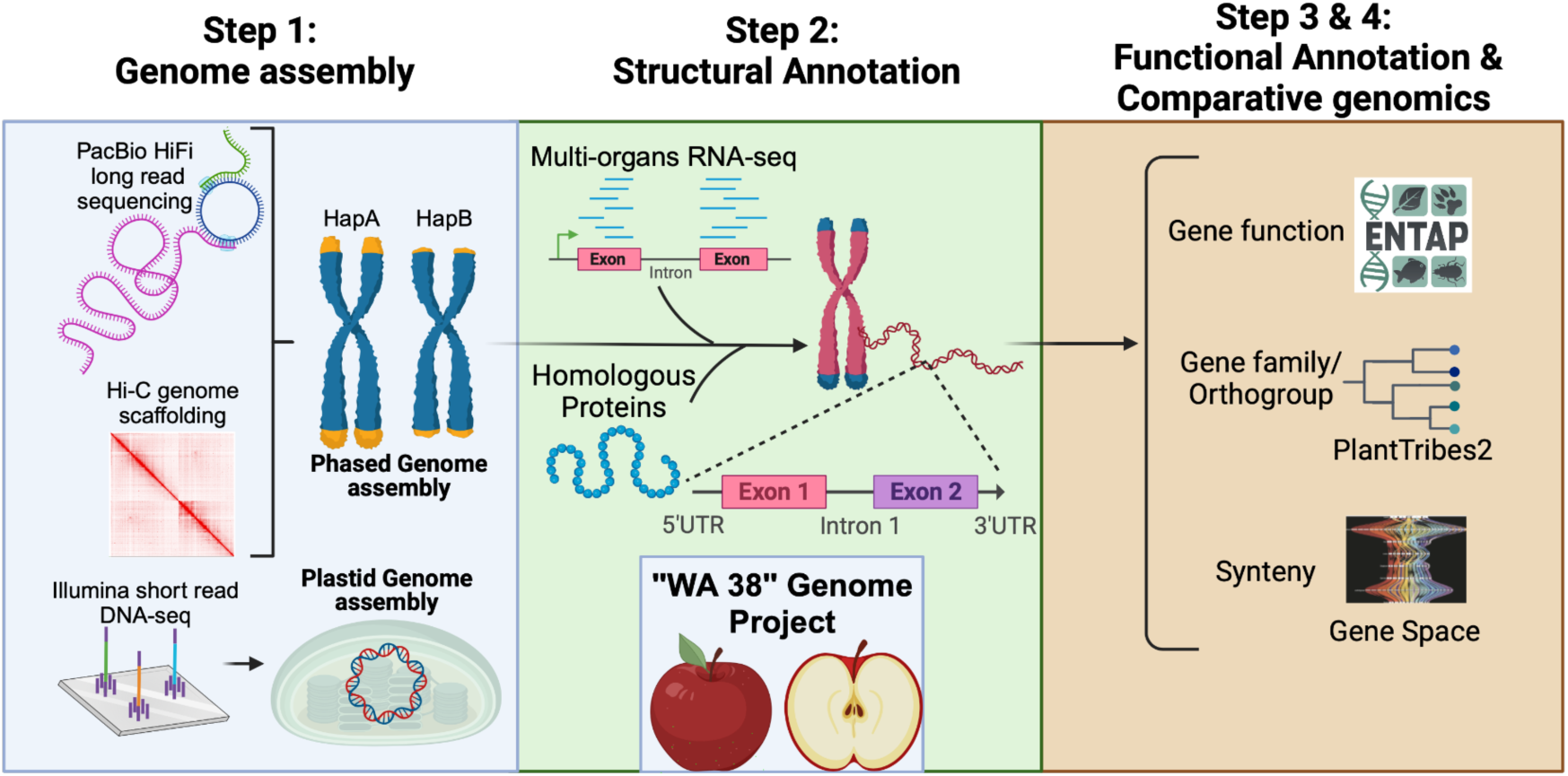
Schematic chart of ‘WA 38’ genome project.

## Methods

Workflows developed for each stage of the project and the summary workflow of the whole project are available in Supplemental Figure S1. Scripts with parameters for each computation step and methods in markdown format are available in GitLab at: https://gitlab.com/ficklinlab-public/wa-38-genome.

### Sample Collection

Approximately 20 grams of young leaf material was harvested from the *Malus domestica* ‘WA 38’ mother tree at the Washington State University and USDA-ARS Columbia View Research Orchard near Orondo, WA, USA and flash-frozen in liquid nitrogen. Tissue was sent to the HudsonAlpha Institute for Biotechnology in Huntsville, AL, USA for DNA extraction, sequencing library preparation, and sequencing, following the same protocol (detailed below) used to generate the ‘d’Anjou’ pear genome (Yocca *et al*. 2024).

To assess heterozygosity and genome size of ‘WA 38’, DNA was extracted using a standard CTAB isolation method (Doyle and Doyle 1987). Illumina TruSeq DNA PCR-free libraries were constructed from 3 ug of input DNA following the manufacturer’s instruction, and sequenced on an Illumina NovaSeq6000.

For PacBio HiFi sequencing, high molecular weight DNA was isolated using a Nanobind Plant Nuclei Big DNA kit (Circulomics-PacBio, Menlo Park, CA), with 4 g of input tissue and a 2-hour lysis. DNA purity, quantity, and fragment sizes were measured via spectrophotometry, Qubit™ dsDNA Broad Range assay (Invitrogen™), and Femto Pulse system (Agilent, Santa Clara, CA), respectively. DNA that passed quality control was sheared with a Megaruptor (Diagenode, Denville, NJ) and size-selected to roughly 25 kb on a BluePippin. The SMRTbell Express Template Prep Kit 2.0 (PacBio, Menlo Park, CA) was used to construct the PacBio sequencing library, and HiFi reads were produced using circular consensus sequencing (CCS) mode with two 8M flow cells on a PacBio Sequel II long-read system.

To scaffold PacBio HiFi contigs into chromosome pseudomolecules, a Dovetail Genomics Omni-C library was generated using 1 g of flash-frozen young leaf material as input following the manufacturer’s instruction (Dovetail Genomics, Scotts Valley, CA), and sequenced on an Illumina NovaSeq6000 S4 PE150 flow cell.

### Sequence quality assessment and genome complexity analysis

Adapter sequences were trimmed from the raw Illumina shot-gun DNA reads using fastp (v0.23.2) (Chen *et al*. 2018) with all the other trimming functions disabled. Both the raw and trimmed Illumina reads, PacBio HiFi reads, and Omni-C Illumina reads were assessed for quality with FastQC (v0.11.9) (Andrews, 2010). Genome complexity, *i.e.* nuclear genome size and ploidy, was estimated using Jellyfish (v2.2.10) (Marçais and Kingsford 2011). With trimmed paired-end Illumina reads as input and a k-mer size set to 21, a k-mer count file was generated by Jellyfish. The k-mer histogram, also created by Jellyfish, was visualized in GenomeScope (v1.0) (Vurture *et al*. 2017) with the following parameters: k-mer size = 21, Read length = 151, and Max k-mer coverage = 1000. A summary statistic report of the sequence quality and complexity analysis was generated with MultiQC (v1.13a).

### Genome Assembly

#### Genome assembly and scaffolding

Phased haplomes were assembled by Hifiasm (v0.16.1) (Cheng *et al*. 2021) with default parameters, using both the Omni-C data and the PacBio HiFi long reads. The statistical summary of the assembly was produced using the assimilation_stats Perl scripts described in (Earl *et al*. 2011). Both hifiasm-assembled haplotype unitigs were then sorted by MUMmer (v3.23) (Kurtz *et al*. 2004) using the ‘nucmer’ function with flag -maxmatch. The resulting files were uploaded to the Assemblytics web server (http://assemblytics.com; Nattestad and Schatz 2016) to visualize structural variations in two haplotype unitigs with the default settings.

Following the initial assembly step, bwa (Li and Durbin 2009) was used to index the draft contigs, and subsequently to align the Hi-C reads to the indexed contigs. The sorted files were input into Phase Genomics hic_qc (https://github.com/phasegenomics/hic_qc; phasegenomics, n.d.) to validate the overall quality of the library. Both assembled haplomes were scaffolded into chromosomes with YaHS (Danecek *et al*. 2021; Zhou *et al*. 2022), using default parameters.

#### Assembly curation, completeness assessment, and telomere identification

Hi-C files were generated using YaHS Juicer Pre (v1.2a.2-0) with flag -a allowing manual curation. The resulting files were used as input for Juicer Tools Pre (v 1.22.01) to generate Hi-C contact maps (Durand *et al*. 2016; Zhou *et al*. 2022). Juicebox Assembly Tools (v1.11.08) was used to explore the Hi-C maps for miss-assemblies (Robinson *et al*. 2018). After manual examination of the Hi-C maps, the final genome assembly was generated by linking remaining files from YaHS Juicer Pre and original HiFi scaffold, using YaHS Juicer Post (v 1.2a.2-0, (Durand *et al*. 2016)).

For consistency and reproducibility, ‘WA 38’ chromosomes were renamed and reorientated to match published genomes. First, MUMmer (v3.23) was used to align the ‘WA 38’ assembly to the ‘Gala’ v1 HapA assembly using the –maxmatch parameter for unique matches (Kurtz *et al*. 2004; Sun *et al*. 2020). Next, Assemblytics dotplot was used to identify ‘WA 38’ scaffolds that aligned with the ‘Gala’ v1 chromosomes and ‘WA 38’ scaffolds were renamed accordingly. To determine orientation, each ‘WA 38’ chromosome was aligned to the corresponding ‘Gala’ v1 chromosomes using LASTZ (v 1.02.00) implemented in Geneious (v9.0.5; Harris. 2007) with the ‘search both strands’ option. The chromosomes on the reverse stand were reoriented with the Reverse Complement (RC) function in Geneious (Supplemental Figure S2). The resulting assembly was searched against NCBI’s RefSeq Plastid database (NCBI Organelle genome resources, n.d.) using megablast and a custom virus and bacteria database using Kraken (v2.1.3; Wood and Salzberg, 2014) to identify contaminants. Scaffolds identified as plastid or microbe contaminants were removed in the assembly.

The cleaned assembly was compared to the ‘Honeycrisp’ genome assembly with a kmer approach using meryl (v1.4.1, Rhie et al., 2020). Chromosomes with a ‘Honeycrisp’ origin were placed in HapA, whereas the others were placed in HapB.

The two final haplome assemblies were compared to each other using MUMmer and Assemblytics as described above to identify structural variants. Benchmarking universal single-copy gene orthologs (BUSCO, v5.4.3_cv1) analysis was performed in genome mode with the eudictos_odb10 database to assess completeness (Manni *et al*. 2021).

### Structural and functional annotation

#### Repeat annotation

Repetitive elements from both haplotypes were annotated using EDTA (v2.0.0; Ou *et al*. 2019), with flags ‘sensitive=1’ and ‘anno=1’. The full coding sequence from ‘Gala’ HapA, obtained from the Genome Database for Rosaceae (GDR; Jung *et al*. 2019), was used as reference to aid repeat finding. The custom transposable element library generated by EDTA was then imported to RepeatMasker (Smit *et al*. 2013-2015) to further identify potentially overlooked repetitive elements and create masked versions of the genome. Three masked versions were generated: softmasked, N masked, and X masked.

Telomeres were identified by tidk (v0.2.41; Brown *et al*. 2023) with the following parameters: explore --minimum 2 --maximum 20 and the default database provided by the software.

#### Gene Annotation

To annotate gene space, a combination of *ab initio* prediction and evidence-based prediction were performed on the softmasked assemblies with two rounds of BRAKER using transcriptome and homologous protein evidence. PASA (v2.5.2; Haas *et al*. 2003) was then used to refine gene models and add UTR annotation. Lastly, a custom script was used for filtering. The detailed methods are described below.

##### BRAKER1 - annotation with transcriptome evidence

To perform transcriptome guided annotation, same RNA-seq data from Khan et al. (2022) (eight tissue types from six pome fruit cultivars including ‘WA 38’, BioProject: PRJNA791346) were first aligned to the ‘WA 38’ haplomes using the STAR aligner implemented in GEMmaker (v2.1.0) Nextflow workflow (Hadish *et al*. 2022). The resulting read alignments were used as extrinsic evidence in BRAKER1 (Hoff *et al*. 2016) to predict gene models in each softmasked haplome with the following parameters: --softmasking, --UTR=off, --species=malus_domestica.

##### BRAKER2 -annotation with homologous protein evidence

To provide protein evidence data for BRAKER2 (Brůna *et al*. 2021), protein sequences from three sources were used: 1) Predicted protein sequences of 13 Rosaceae genomes retrieved from GDR (*Fragaria vesca* v4a2 (Li *et al*. 2019), *Malus baccata* v1.0 (Chen *et al*. 2019), *M. domestica* var.Gala v1 (Sun *et al*. 2020), *M. domestica* var.GDDH13 v1.1 (Daccord *et al*. 2017), *M. domestica* ‘Honeycrisp’ v1.0 (Khan *et al*. 2022), *M. sieversii* v1 (Sun *et al*. 2020), *M. sylvestris* v1 (Sun *et al*. 2020), *Prunus persica* v2.0.a1 (Verde *et al*. 2017), *Pyrus betulifolia v1.0 (Dong et al. 2020)*, *P. communis* ‘d’Anjou’ v2.3 (Yocca *et al*. 2023), *P. pyrifolia* ‘Nijisseikiv’ v1.0 (Shirasawa *et al*. 2021)*, Rosa chinensis* ‘Old Blush’ v2.0.a1 (Raymond *et al*. 2018), and *Rubus occidentalis* v3 (VanBuren *et al*. 2018)); 2) Peptide sequences predicted from *de novo* transcriptome assemblies used in the ‘Honeycrisp’ genome annotation (Khan *et al*. 2022); and 3) Viridiplantae OrthoDBv11 protein sequences (Kuznetsov *et al*. 2022). In the same manner as BRAKER1, the softmasked haplome assemblies were used as input.

##### TSEBRA - transcript selection

The gene annotation results from BRAKER1 and BRAKER2 were merged and filtered based on the supporting evidence using TSEBRA (v.1.0.3; Gabriel *et al*. 2021) with the default configuration (file obtained in August 2022) provided by TSEBRA developers.

##### PASA - gene model curation and UTR annotation

Two sources of transcriptome assembly evidence were obtained to facilitate PASA annotation: 1) Transcript sequences predicted from *de novo* transcriptome assemblies used by ‘Honeycrisp’ genome annotation; and 2) Reference guided assemblies created with read alignment files from GEMmaker (see the BRAKER1 section for details) using Trinity (Grabherr et al. 2011) with max intron size set to 10,000. Four rounds of PASA (v2.5.2) curation were performed using the aforementioned evidence and a starting annotation. The first round of PASA curation used TSEBRA annotation as the starting annotation, and annotations from the previous round were used as the starting annotation for rounds two through four. The curation results from each round were manually inspected using the PASA web portal. No significant improvement was observed after the fourth round of curation, therefore no further rounds were performed.

##### Gene model filtering and gene renaming

Repeat and gene model annotations were loaded to IGV (v2.15.1; Robinson *et al*. 2011) for manual inspection. Three types of erroneous gene models were observed consistently throughout the annotations. Type 1: Genes overlapping with repeat regions (e.g. transposon was wrongly annotated as a gene), Type 2: Gene models overlapping with each other on the same strand (e.g. single gene was wrongly annotated with multiple gene models), and Type 3: Gene models with splice variants that had no overlap (e.g. different genes were wrongly annotated as the single gene’s splice variants). A custom script was used to address these errors. The Type 1 error was resolved by removing genes with 90% of its coding region overlapping with repeat regions. The Type 2 error was resolved by removing the shorter gene of a pair that overlaps on the same strand. The Type 3 error was resolved by splitting splice variant models with no overlap into two separate gene models. Finally, custom scripts were used to generate the final annotation files (gene, mRNA, cds, protein, gff3) and rename genes to match the naming convention proposed by GDR (https://www.rosaceae.org/nomenclature/genome). The longest isoforms of each transcript were needed for some downstream analysis and were extracted using a modified version of the get_longest_isoform_seq_per_trinity_gene.pl script provided by Trinity (Grabherr *et al*. 2011).

#### Functional Annotation

The final gene sets from both ‘WA 38’ haplomes were annotated using EnTAPnf (Hart *et al*. 2020) with Interproscan, Panther, RefSeq, and uniprot_sprot databases that are automatically downloaded using the download.py script provided by EnTAPnf.

### Comparative Analysis

#### Synteny analysis

A synteny comparison was performed using GENESPACE (Lovell *et al*. 2022) with five *Malus domestica* assemblies and annotations (GDDH13 from Daccord *et al*. 2017), both haplomes of ‘Honeycrisp’ from Khan *et al*. 2022, and both haplomes from ‘WA 38’). Default parameters were used. Only the longest isoforms were used for ‘WA 38’.

#### Gene family analysis

Gene family, or orthogroup, analyses were carried out to identify shared and unique gene families in ‘WA 38’ and other pome fruit genomes (i.e., *Malus* sp. and *Pyrus* sp. A full list of genomes analyzed can be found in Supplemental Table S1) following the method described by (Khan *et al*. 2022). Briefly, predicted protein sequences from the selected pome fruit genomes were classified into a pre-computed orthogroup database (26Gv2.0) using the ‘both HMMscan and BLASTp’ option implemented in the GeneFamilyClassifier tool from PlantTribes2 (Wafula *et al*. 2022). Overlapping orthogroups among *M. domestica* genomes were calculated and visualized with the UpSet plot function implemented in TBtools v2.030 (Chen *et al*., 2023).

A Core OrthoGroup (CROG) - Rosaceae gene count analysis was carried out following the method described by (Wafula *et al*. 2022). First, a CROG gene count matrix was created by counting genes classified into CROGs from each pome fruit genome. Next, the matrix was visualized as a clustermap using the Seaborn clustermap package (CROGs with standard deviation of 0 were removed prior to plotting) with rows normalized by *z*-score. Finally, the derived *z*-score of CROGs in each genome was summarized into a boxplot to illustrate z-score distribution using the boxplot function in Seaborn.

### Gene evidence source mapping

Each gene was screened against the following evidence source: Transcriptome evidence covering the entire gene (Full support); Transcriptome evidence covering part of the gene (Any support); Homologous protein evidence covering the entire gene (Full support); Homologous protein evidence covering part of the gene (Any support); Has a EnTAP functional annotation from any database; Assignment to a PlantTribes2 Orthogroup. Transcriptome and homologous protein evidence were mapped to genes by using “selectSupportedSubsets.py” script provided by BRAKER (Brůna *et al*. 2021) and BEDtools (Quinlan and Hall 2010). Summaries of evidence source mapping are available in Supplemental Table S2 and S3. The following subsets of genes were extracted and were subject to BUSCO completeness analysis and CROG gene count analysis: Subset 1, Genes with ***full*** support from either RNAseq or homologous protein evidence; Subset 2, Genes with ***any*** support from either RNAseq or homologous protein evidence; Subset 3, Genes from Subset 1 plus gene with both EnTAP and PlantTribes2 annotation; Subset 4, Genes from Subset 1 plus genes with either EnTAP or PlantTribes2 annotation.

### Chloroplast & Mitochondria Assembly and Annotation

The chloroplast genome was assembled from trimmed Illumina shot-gun DNA reads using NOVOplasty (v4.3.1; Dierckxsens *et al*. 2017) with the *Malus sierversii* chloroplast genome (NCBI accession ID: MH890570.1; Naizaier *et al*. 2019) as the reference sequence and the NOVOplasty *Zea mays RUBP* gene as the seed sequence. The assembled chloroplast was annotated using GeSeq Web Server (website accessed on Dec. 19th, 2023; Tillich *et al*. 2017) with settings for ‘circular plastid genomes for land plants’ and the following parameters: annotating plastid inverted repeats and plastid trans-spliced *rps12*. Additionally, annotations from third party softwares Chloё (v0.1.0) and ARAGORN (v1.2.38), as well as a BLAT (v.35×1) search against all land plant chloroplast reference sequences (CDS and rRNA), were integrated with the GeSeq results. Genes identified by multiple tools were manually reviewed to produce the final, curated annotation. The curated chloroplast annotation was visualized by OGDRAW (v1.3.1; Greiner *et al*. 2019).

The mitochondrial genome sequence was isolated from the Hifiasm assembled contigs using MitoHifi (v3.2; Uliano-Silva *et al*. 2023). The *M. domestica* mitochondria sequencing from NCBI (NC_018554.1; Goremykin *et al*. 2012), which contained 57 genes consisting of 4 rRNAs, 20 tRNAS, and 33 protein-coding genes, was used as the closely related reference sequence. Briefly, MitoHifi compares the assembled contigs to the reference mitogenome using the BLAST algorithm. The resulting contigs were manually filtered by size and redundancy and then are circulated. To increase the annotation quality, GeSeq was deployed in mitochondrial mode with the *M. domestica* NCBI RefsSeq sequence to annotate the ‘WA 38’ mitochondria assembly. Fragmented genes from the annotation were manually removed prior to visualization in OGDRAW (v1.3.1; Greiner *et al*. 2019).

## Results

### A Complete, Reproducible, Publicly-available Workflow

To ensure transparency and reproducibility, the ‘WA 38’ Whole Genome Assembly and Annotation (WA 38 WGAA) project workflow was made publicly accessible through a GitLab repository (https://gitlab.com/ficklinlab-public/wa-38-genome). This repository contains the complete manual workflow for assembly and annotation of the genome. It organizes each step in order of execution, using ordered, numeric directory prefixes where each directory includes detailed method documentation and scripts that were executed for each analysis. All parameter settings, as well as any command line manipulation of the files generated are noted in the scripts or methods. Summary diagrams for the manually executed workflow are available in Supplemental Figure S1. All software utilized in the project has been containerized and shared on Docker Hub (https://hub.docker.com/u/systemsgenetics). Any user that follows the workflow can retrieve the public data and repeat the steps to reproduce the results. Leveraging these resources from the ‘WA 38’ WGAA project, and as part of our commitment to knowledge sharing, we have initiated an American Campus Tree Genome (ACTG) course GitHub organization (https://github.com/actg-course/). This organization comprises three main repositories: 1) wgaa-compute: a generic whole genome assembly and annotation workflow template, derived from the ‘WA 38’ WGAA project, that can be adapted for other species; 2) wgaa-docker: the Docker recipes for all the software employed in the project; and 3) wgaa-doc: an open-source and editable documentation repository containing teaching materials for current and future ACTG instructors, providing a collaborative space for instructors to learn from and contribute to the enhancement of the course materials.

### Nuclear Genome Assembly

#### Sequence Quality Assessment

Raw sequencing data (Table 1) was assessed for read quality. The Illumina shotgun short read data consisted of 807.2 million total reads with a mean length of 151bp for a total of 121.9 Gigabases (Gb) of data after adapter trimming. Filtered Illumina data GC content is 38% and has 91.8% Q20 bases and 83.4% Q30 bases. Duplication rates ranged from 23.3% to 27.8%. PacBio long read raw data consisted of 3.9 million reads from 85-49,566bp in length for a total of 60.0 GB. Sequence duplication rates ranged from 2.2% to 2.4%. PacBio sequence GC content is 38%, same as the Illumina data. In addition, a 402x coverage (201x for each haplome) of Omni-C data was generated to facilitate the assembly and phasing.

**Table 1.** Yield of Illumina DNA short reads (Shotgun and Omni-C) and PacBio HiFi sequencing reads from young leaf tissues of ‘WA 38’.

|  | Long Read | Short Read |  |
| --- | --- | --- | --- |
|  | PacBio HiFi | Shotgun DNA seq | OmniC-Seq |
| Total read number | 3,870,263 | 807,220,896 | 1,730,268,360 |
| Number of bases (Gb) | 60.0 | 121.9 | 261.3 |
| Coverage* | 92x | 188x | 402x |
| Average length (bp) | 15,495 | 151 | 151 |

\* calculated with the size of a haploid genome (650 Mb).

#### Genome complexity

Using a *k*-mer frequency approach, genome characteristics such as heterozygosity and genome size were estimated (Figure 3). Analysis of both short and long reads resulted in an estimated heterozygosity of ∼1.35%, similar to estimates from the ‘Honeycrisp’ cultivar (1.27%; Khan *et al*. 2022). Estimate genome size was 467Mb from the short reads and 606Mb from the long reads. These estimates are lower than expected from other apple genomes (‘Honeycrisp’: 660-674 Mb; Khan *et al*. 2022 and ‘Golden Delicious’: ∼701 Mb; Li *et al*. 2016) and the final assembly (Table 2). Additionally, the percent of unique sequence was estimated at 69.5% for the short reads and 53.4% for the long reads, with the long read estimate being more consistent with what is expected from the ‘Honeycrisp’ (51.7%; Khan *et al*. 2022) and of wild apple species *Malus baccata* (58.6%; Chen *et al*. 2019).

**Figure 3.**
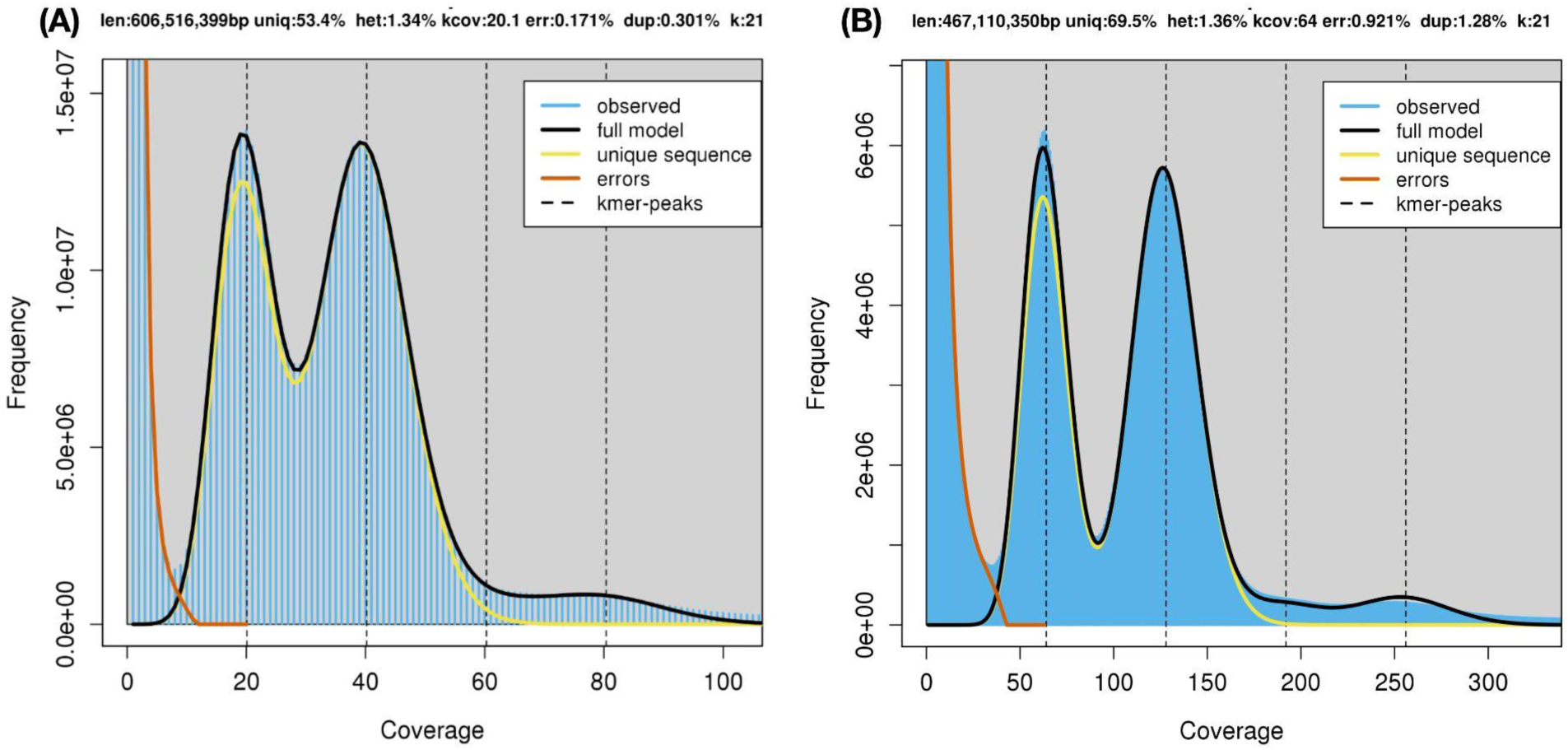
Genome complexity of ‘WA 38’ genome using PacBio long read data (A) and Illumina short read (B). The output figure was generated by GenomeScope (k=21).

**Table 2.** Comparison of genomic features and assembly statistics of the ‘WA 38’ genome and previously published apple genomes.

|  | ‘WA 38’ |  | ‘Honeycrisp’ |  | ‘Antonovka’ | ‘Gala’ | GDDH13 | ‘Fuji’ |
| --- | --- | --- | --- | --- | --- | --- | --- | --- |
|  | HapA | HapB | HapA | HapB |  |  |  |  |
| Number of Scaffold | 17 | 17 | 473 | 215 | 168 | 812 | 1,081 | 1,358 |
| Haploid genome size (Mb) | 645.41 | 651.07 | 674 | 660 | 643.5 | 652.4 | 709.6 | 736.9 |
| N50 (Mb) | 36.1 | 37.2 | 31.6 | 32.8 | 35.85 | 23.9 | 5.5 | 36.8 |
| L50 | 8 | 8 | 8 | 8 | 8 | 8 | NA | 9 |
| Number of protein-coding genes | 53,028 | 54,235 | 47,563 | 48,655 | 45,085 | 45,352 | 45,116 | 49,972 |
| Complete BUSCO (%) Assembly | 98.7 | 98.7 | 98.6 | 98.7 | 97.6 | 97.9 | 98 | 98.8 |
| Complete BUSCO (%) Annotation | 98.5 | 98.4 | 96.8 | 97.4 | 97.25 | 95.5 | 96.1 | 97.2 |
| Number of orthogroups in 26Gv2.0 | 10,494 | 10,511 | 10,350 | 10,366 | 10,293 | 10,095 | 10,117 | 10,243 |
| Reference | This paper |  | Khan et al. 2022 |  | Švara et al., 2023 | Sun et al. 2021 | Daccord et al. 2017 | Li et al., 2024 |

*NA: Data not available. For consistency, genome statistics and BUSCO analyses were performed on the publicly available genomes using the same methods used for ‘WA 38’, except for N50 and L50 of GDDH13 as the scaffold assembly is not publicly available. ‘Antonovka’ data is the average of the two haplomes. The unphased version of ‘Fuji’ was used. A more in-depth comparison is available in Supplemental Table S5.*

#### Genome assembly, scaffolding, and curation

For initial assembly, scaffolding, and curation, two unsorted, phased haplomes, called Hap1 and Hap2, were assembled and scaffolded using both PacBio long reads and Omni-C short reads. Hi-C maps of the haplome assemblies show no mis-assemblies (Supplemental Figure S3). For Hap1 and Hap2, a total of 22 joins and 20 joins, respectively, were made in the scaffolding step to build the final assemblies into 17 chromosomes each, with the remaining scaffolds representing unincorporated contigs. Unincorporated contigs were investigated and found to be bacterial or other contamination and were removed. After removing contaminants, Hap1 is 645.41 Mb in length with an N50 of 36.1 Mb, while Hap2 is 651.07 Mb in length with an N50 of 37.2 Mb. Additional assembly statistics for both haplomes are included in Supplemental Table S4. ‘WA 38’ has a comparable genome size to other previously sequenced apple cultivars, including its parent ‘Honeycrisp’ (Khan *et al*. 2022). Notably, the ‘WA 38’ scaffold N50 is among the longest across all published apple genomes, indicating high levels of assembly contiguity (Supplemental Table S5).

#### Haplotype-binning, Structural Comparison, and Completeness Assessment

The K-mer based binning method identified the origin of chromosomes in each haplome assembly. Ten out of the 17 chromosomes in Hap1 originated from ‘Honeycrisp’, while the other seven were from ‘Enterprise’. After reorganizing the chromosomes based on parent contribution, the haplome containing all the ‘Honeycrisp’ origin chromosomes is designated as HapA, whereas the ‘Enterprise’ originated haplome is designated as HapB. HapA and HapB are structurally similar; a total of ∼44 Mb are affected by structural variants and are mainly contributed by indels and repeat expansion and contractions (Supplemental Table S6 and Supplemental Figure S4). Additionally, three large inversions are observed on chromosomes 1, 11, and 13 (Supplemental Figure S5). Based on the BUSCO analysis, both the HapA and HapB assemblies were 98.7% complete, with only 19 BUSCOs missing and 12 partially detected (Supplemental Table S7). This BUSCO score suggests high genome completeness for both haplomes, comparable to the ‘Fuji’ apple genome assemblies, which is most contiguous of all apple genomes to date (Table 2 and Supplemental Table S5; Li et al., 2024).

### Nuclear Genome Structural Annotation

#### Repeat annotation

In both haplomes, approximately 58.7% of the assembly was predicted to be repetitive regions by EDTA (Ou *et al*. 2019; Table 3). RepeatMasker identified an additional 4% repeat elements, resulting in a total of 62.7% repeat regions in both HapA and HapB, comparable to the ‘Honeycrisp’ genome (Khan *et al*. 2022). In both haplomes, the most dominant type of repeat element is long terminal repeat (LTR), followed by Terminal Inverted Repeat (TIR) (Table 3, Supplemental Table S8), consistent with that in ‘Honeycrisp’. We also compared the repeat landscape of ‘WA 38’ with ‘d’Anjou’ pear which was annotated with the same methodology. While they share the major repeat classes, ‘d’Anjou’ pear has a much lower percentage of repeat elements (Table 3).

**Table 3.** Summary of repetitive element annotation in the ‘WA 38’ and other apple genomes.

| Class |  | ‘WA 38’ (%) |  | ‘Honeycrisp’ (%) |  | ‘d’Anjou’ pear (%) |  |
| --- | --- | --- | --- | --- | --- | --- | --- |
|  |  | HapA | HapB | HapA | HapB | Hap1 | Hap2 |
| LTR | Copia | 9.37 | 10.22 | 9.73 | 9.6 | 5.6 | 5.73 |
|  | Gypsy | 17.19 | 18.32 | 20.29 | 17.8 | 12.32 | 12.88 |
|  | unknown | 16.37 | 14.52 | 14.89 | 16.86 | 8.46 | 10 |
| TIR | CACTA | 1.94 | 2.15 | 2.21 | 1.95 | 1.4 | 1.4 |
|  | Mutator | 3.96 | 4.18 | 4.16 | 4.25 | 3.47 | 3.41 |
|  | PIF Harbinger | 2.4 | 2.52 | 2.43 | 2.6 | 1.81 | 1.81 |
|  | Tc1_Mariner | 0.16 | 0.24 | 0.15 | 0.27 | 0.13 | 0.11 |
|  | hAT | 2.15 | 2.37 | 2.3 | 2.31 | 0.58 | 0.84 |
|  | polinton | 0 | 0 | 0 | 0.01 | 0 | 0 |
| nonLTR | LINE_element | 0.14 | 0.16 | 0.18 | 0.17 | 0.14 | 0.14 |
|  | unknown | 0.11 | 0.09 | 0.09 | 0.18 | 0.06 | 0.06 |
| nonTIR | helitron | 3.41 | 2.20 | 2.95 | 3.18 | 1.56 | 1.92 |
| Other repeat region |  | 1.52 | 1.74 | 2.91 | 2.78 | 3.98 | 4.22 |
| RM* |  | 3.98 | 3.99 | NA | NA | NA | NA |
| Total |  | 62.71 | 62.71 | 62.43 | 61.97 | 39.78 | 42.52 |
| Reference |  | This paper |  | Khan et al. 2022 |  | Yocca et al., 2023 |  |

\* repeat regions annotated by RepeatMakser

Through telomere search in each haplotype, we discover that telomere repeat regions are present in almost every chromosome of each haplome. The most enriched telomere repeat unit is a 7-mer “AAACCCT” and its reverse complement “AGGGTTT”, which has been reported as overrepresented in the *Arabidopsis thaliana* genome (Choi *et al*. 2021), opposed to “CCCATTT” and “TTTTAGGG” reported in the most recent T2T ‘Golden Delicious’ apple genome (Su *et al*. 2024). A list of telomere repeat regions and units for both haplotypes were deposited in Supplemental Table S9.

#### Gene space annotation

To annotate the gene space, we utilized a combination of *ab initio* prediction and evidence-based prediction with transcriptome and homologous protein, functions implemented in BRAKER2 (Brůna *et al*. 2021). However, BRAKER2 was unable to annotate UTR regions and yielded erroneous gene models and splice variants (Supplemental Figure S6). Therefore, the gene models were further processed with PASA (Haas *et al*. 2003) and a custom script. A total of 53,028 and 54,235 genes were annotated from HapA and HapB, respectively, more than most published apple genomes (Table 2, Supplemental Table S10). The complete BUSCO scores for HapA and HapB annotations are 98.5% and 98.4%, respectively, the highest score among all *M. domestica* genomes sequenced to date (Supplemental Table S5). The average protein annotated from HapA and HapB contains 361.3 and 356.4 amino acids, respectively, similar to that of other *M. domestica* annotations (Supplemental Table S11). On average, 1.3 splice variants were identified for each gene in both HapA and HapB annotations. The only other apple genome with splice variant annotation is ‘Honeycrisp’, and on average, 1.05 splice variants were annotated per gene (Supplemental Table S11). Additionally, 53.5% and 52.2% of the annotated transcripts from HapA and HapB, respectively, contain untranslated regions (UTRs). Notably, ‘WA 38’ is the only other apple genome besides ‘GDDH13’ and ‘Fuji’ that has more than half of the genes annotated with UTRs.

The ‘WA 38’ genes were named in accordance with the convention following guidance from the Genome Database for Rosaceae (GDR). This convention was first proposed by our group for the ‘Honeycrisp’ genome and was later adopted with modification by GDR (Gene name example: *drMalDome.wa38.v1a1.ch10A.g00001.t1*). This convention meets recommendations proposed by the AgBioData consortium to reduce gene ID ambiguity and improve reproducibility.

### Nuclear Genome Functional Annotation

EnTAP (Hart *et al*. 2020) functional annotation assigned functional terms to 89.5% and 88.8% of proteins annotated from HapA and HapB, respectively. Specifically, an average of 83% and 55% of all proteins (including both HapA and HapB) have strongly supported hits in the NCBI RefSeq (O’Leary *et al*. 2016) and UniProt database, respectively, 75% were annotated with an InterPro term, and 88% have functional annotations from at least one of the databases included in InterProScan. EggNOG (O’Leary *et al*. 2016; Huerta-Cepas *et al*. 2019) search provided additional function information: 90% of the annotated proteins were assigned into EggNOG orthogroups, 84% were annotated with protein domains, 21% were classified into KEGG pathways, and 63%, 53%, and 61% proteins were annotated with GO biological process, cellular component, and molecular function terms, respectively (Supplemental Table S12).

### Comparative Analyses

Synteny and gene family analyses were performed to investigate the similarity and unique features of ‘WA 38’ genome to other closely related species and cultivars.

Synteny analysis was performed to compare the genomes of ‘WA 38’, one of its parents, ‘Honeycrisp’, and the most referenced apple genome, ‘GDDH13’, using GeneSpace. The two ‘WA 38’ haplomes are highly collinear with each other and with the other apples, especially the two ‘Honeycrisp’ haplomes. Although inversions at various scales were observed between the two ‘WA 38’ haplomes e.g. large inversions on chromosomes 1, 11, 13 (Supplemental Figure S5 and S7), they have minor effects on gene order (Figure 4), likely due to the small number of genes annotated from those inverted regions.

**Figure 4.**
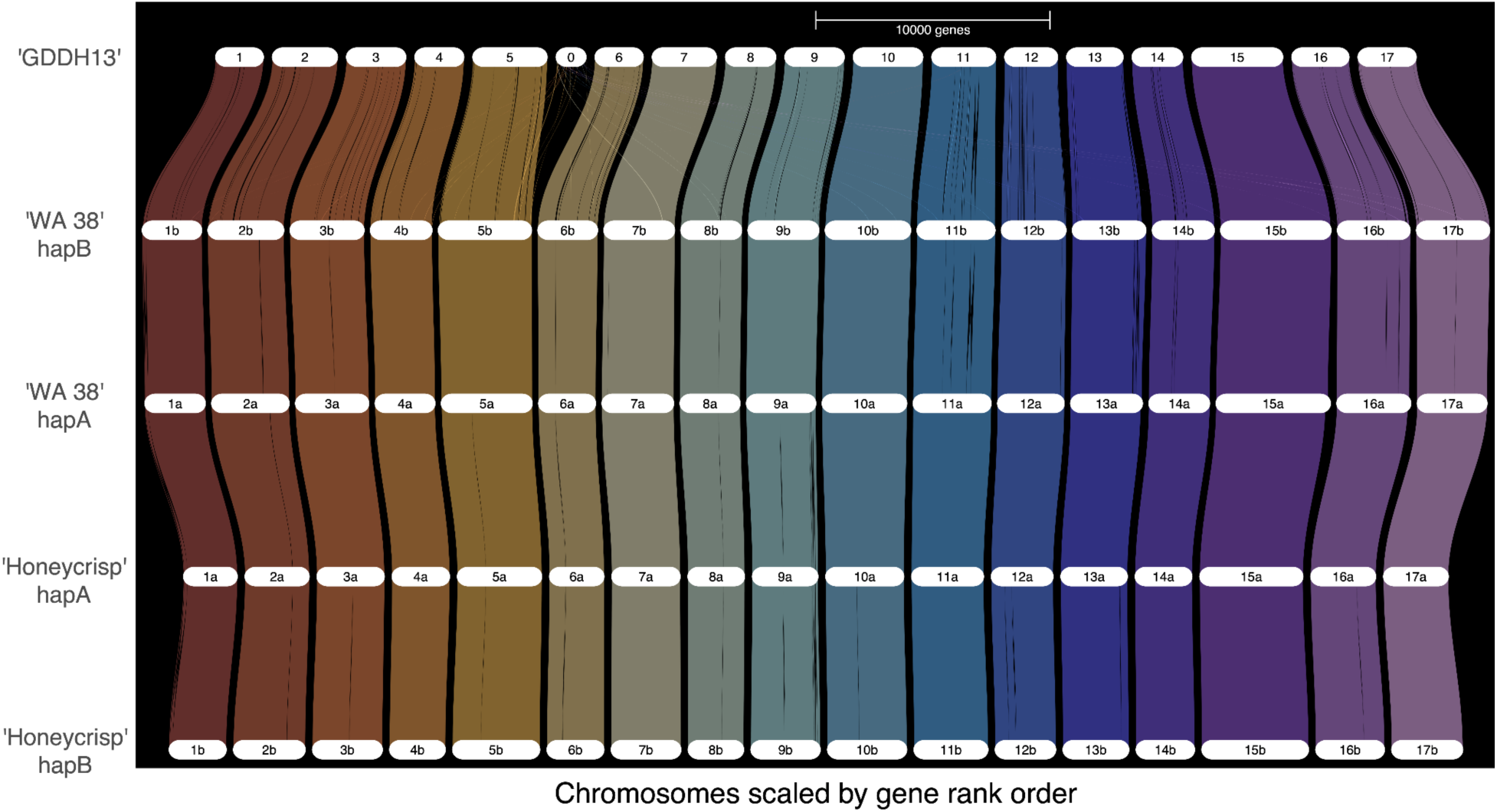
Riparian plot comparing ‘WA 38’ Haplotype A and B with ‘Honeycrisp’ Haplotype A and B and ‘Golden Delicious’ (GDDH13) genomes by gene rank order.

Gene family analysis is performed using PlantTribes2 and the pre-constructed 26Gv2.0 scaffold orthogroup database (Wafula *et al*. 2022). Out of the 18,110 pre-constructed orthogroups, proteins from all apple annotations (including 6 published scion cultivar genomes, 2 rootstock genomes, and the ‘WA 38’ genome from this work) are found in 11,698 orthogroups. ‘Golden Delicious’ Genome v1.0 (Velasco *et al*. 2010) was omitted from this analysis due to poor annotation quality. Proteins from HapA and HapB of ‘WA 38’ were classified into 10,494 and 10,511 orthogroups, respectively, similar or slightly higher in number compared to previously published *M. domestica* genomes, including ‘Honeycrisp’, ‘Gala’, and ‘GDDH13’ (Table 2, Figure 5). An investigation into shared and unique orthogroups across all the scion genomes showed that most orthogroups (8,800 or 75%) are shared by all six apple genomes considered. Additionally, 824 orthogroups are shared by both ‘WA 38’ haplomes and the seven other annotations (each of the two haplomes from ‘Honeycrisp’ and ‘Antonovka 172670-B’ are counted as unique annotations). ‘Honeycrisp’ shared the largest number of orthogroups with ‘WA 38’, as expected due to being a parent of ‘WA 38’ (Supplemental Table S13). These results indicate that the ‘WA 38’ annotation captures genes in virtually all *M. domestica* orthogroups. Additionally, 39 orthogroups were unique to ‘WA 38’ (*i.e.* present only in the two ‘WA 38’ haplomes) and each haplome of ‘WA 38’ contains 44 unique orthogroups (Figure 5).

**Figure 5.**
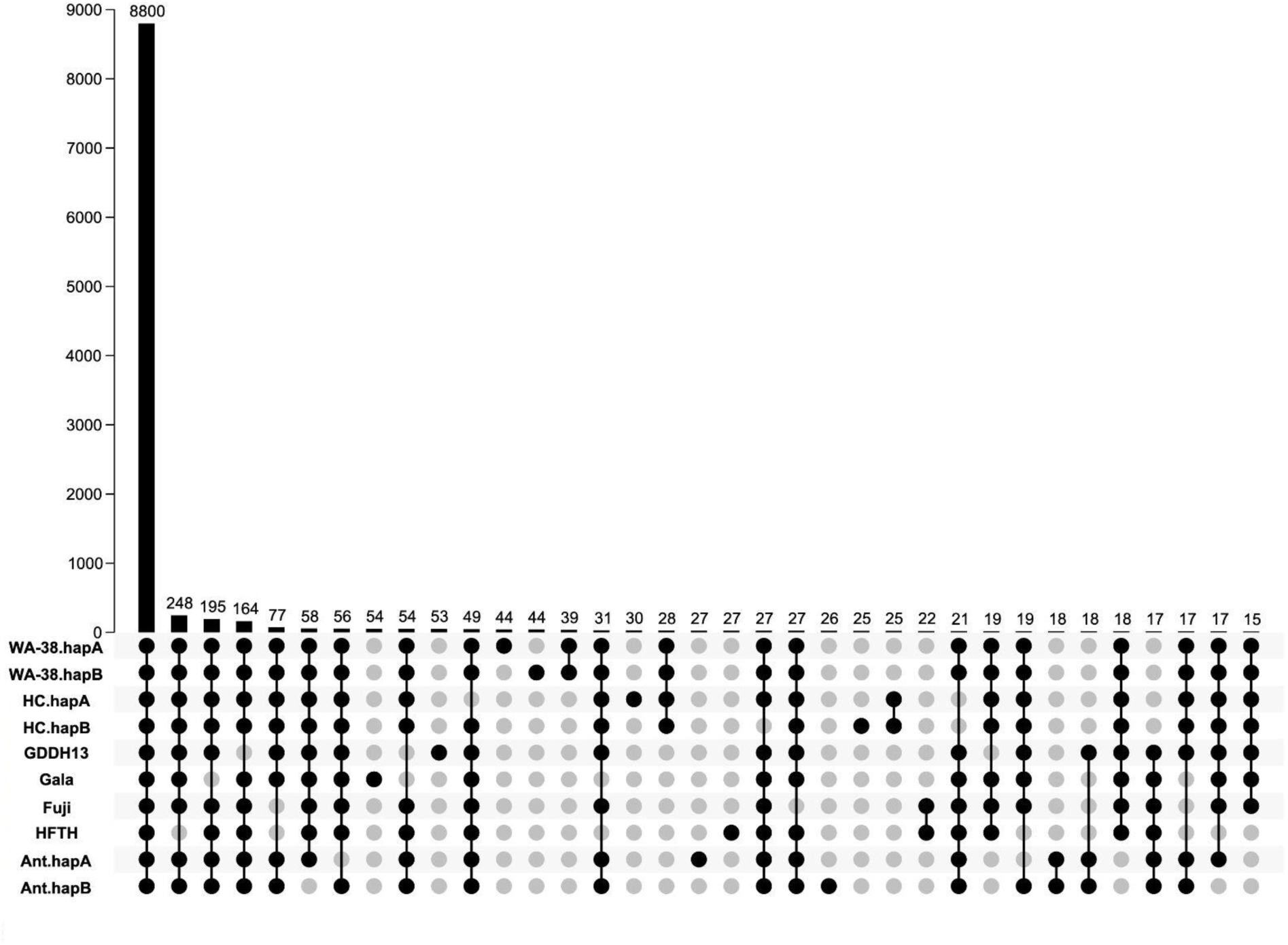
Upset plot of shared and unique orthogroups among Malus domestica genomes. Rows in the bottom of the figure are genomes used for the comparison. Columns (categories, x-axis of the bar graph) are annotated with black or gray dots where black is present and gray is absent. The height of the black bars (y-axis of the bar graph) is scaled to match the number of orthogroup in each category, which are also printed above the bars.

In addition to identifying the shared and unique orthogroup, a CoRe OrthoGroup (CROG) - Rosaceae analysis was performed to further investigate orthogroup contents. As expected, in the CROG gene count clustermap (Figure 6), ‘WA 38’ clustered closely with ‘Honeycrisp’. The ‘WA 38’ + ‘Honeycrisp’ group is clustered with ‘GDDH13’, as expected based on pedigree (Howard *et al*. 2017). Interestingly, a strong ‘publication bias’, first mentioned by Wafula et al., 2022, is observed: genomes released in the same publication or annotated by the same researcher clustered together. Such groups are: ‘Gala’, *Malus sieversii*, and *M. sylvestris* (Sun *et al*., 2021); ‘Fuji’, ‘M9’, and ‘MM106’ (Li *et al*., 2024); *M. fusca* (Mansfeld *et al*. 2023) and *Pyrus communis* ‘d’Anjou’ (Yocca *et al*. 2024); ‘Honeycrisp’ (Khan *et al*. 2022) and ‘WA 38’. The CROG gene count *z*-score box plot shows (Figure 7) that the average *z*-score of ‘WA 38’ gene counts are slightly higher than expected (with 0 as the perfect score), indicating that there are a number of CROGs containing more genes from the ‘WA 38’ annotations compared to other apples.

**Figure 6.**
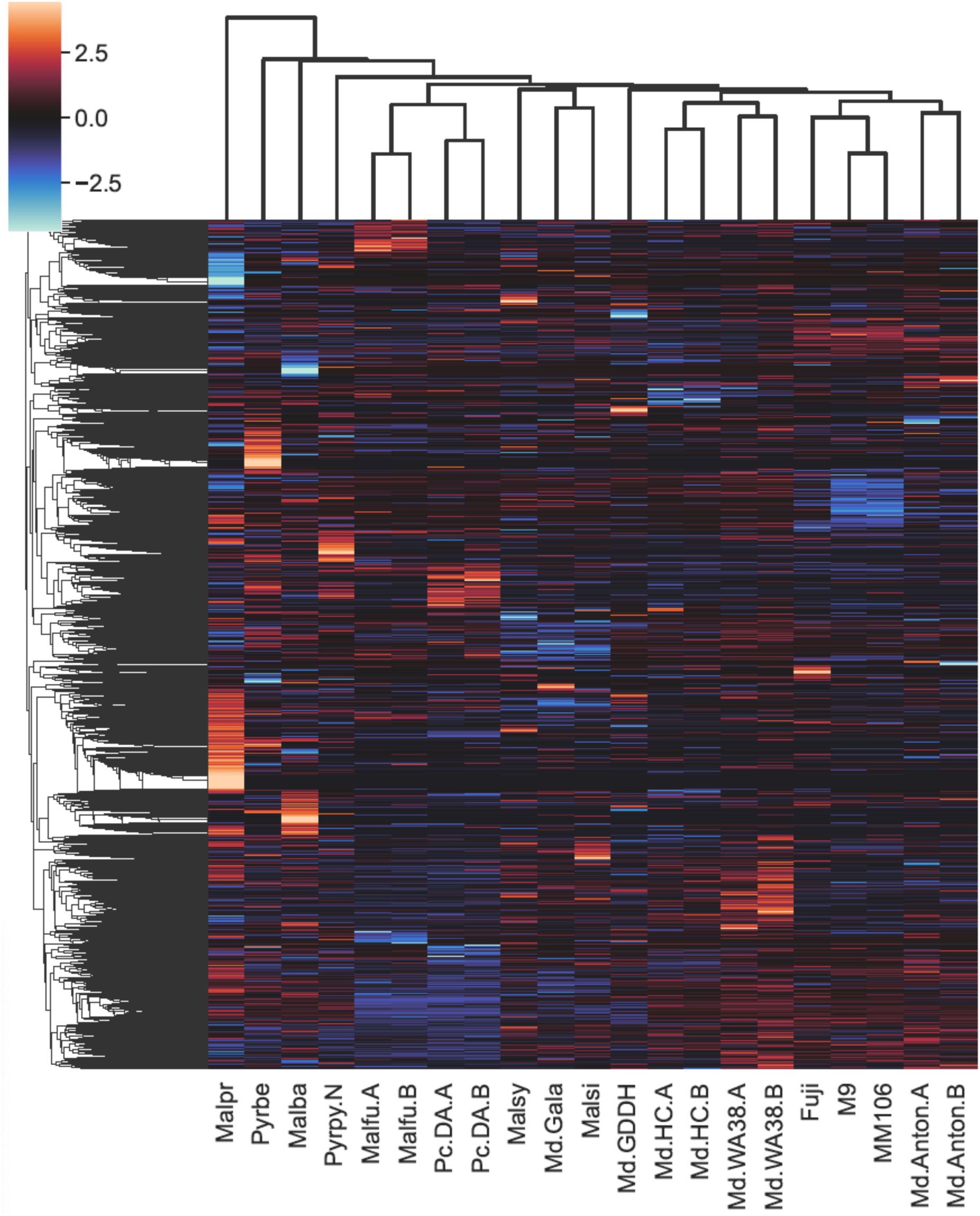
CoRe OrthoGroup (CROG) - Rosaceae gene count clustermap. Each row represents a CROG and each column represents a genomes. Color indicates the number of genes in each cell relative to the row average (z-score). Warmer color indicates more genes. Cooler color indicates fewer genes. The darker a color, the closer the value is to the row average. Genome and annotation abbreviations can be found in Supplemental Table 1.

**Figure 7.**
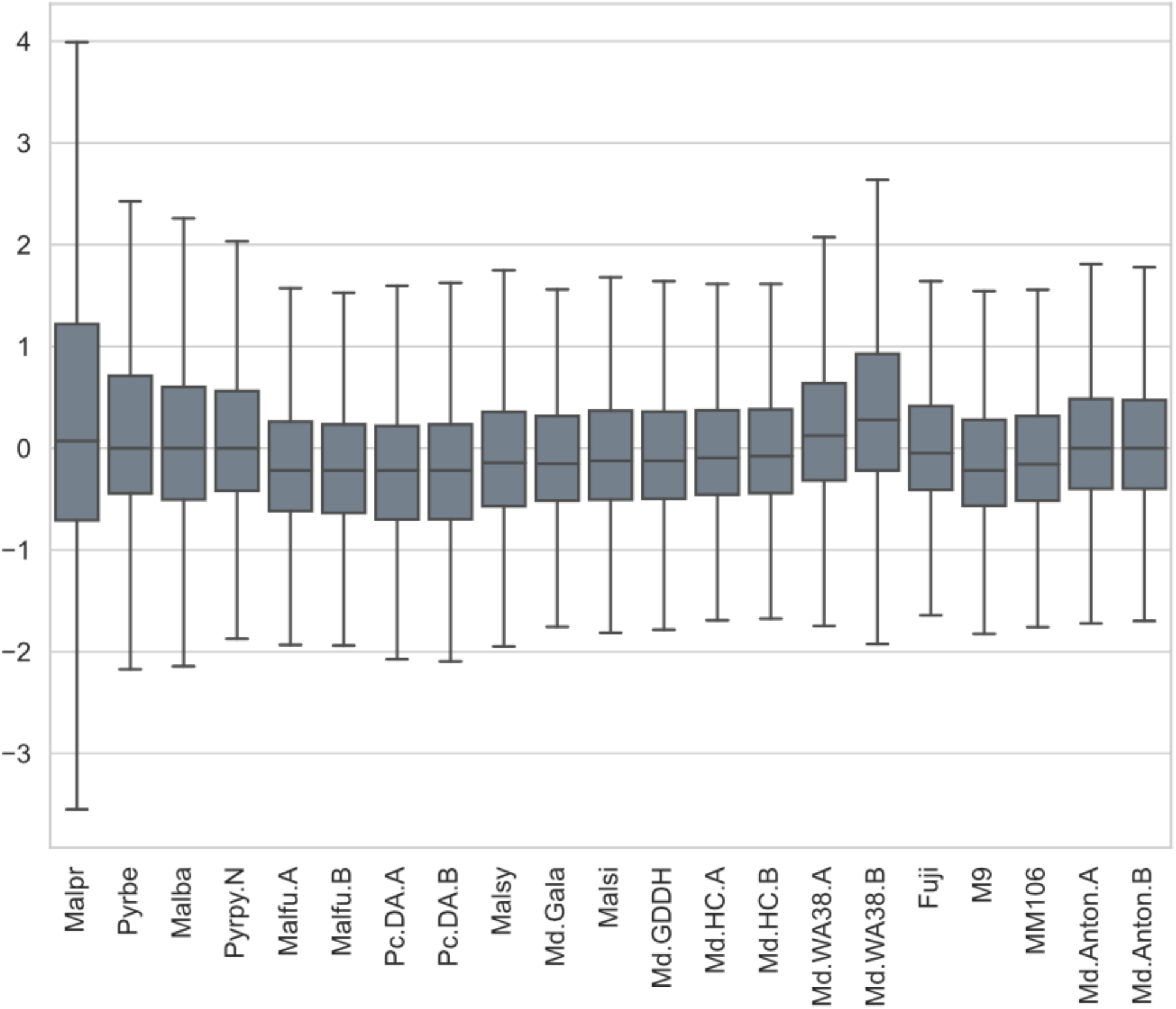
Boxplot summarizing z-score distribution of CROG gene counts in selected pome fruit genomes. Genome and annotation abbreviations can be found in Supplemental Table 1.

### Gene model evidence source mapping

The final gene model annotation contains *ab initio* prediction and genes with transcript evidence and/or homologous protein support. Although high BUSCO completeness scores are obtained from both haplome annotations, their gene numbers are greater than expected (45,000-49,000 based on previous publications). Therefore, we explored evidence supporting a gene model to be a true positive, including extrinsic evidence (from transcript and homologous protein) used in gene model annotation and comparative genomic evidence (EnTAP functional annotation and gene family circumscription), and assessed completeness via a BUSCO analysis (Table 4). The most stringent filter, the same strategy deployed in the ‘Honeycrisp’ genome annotation, was to remove genes without full support from both transcript and homologous protein evidence (Subset 1 in Table 4). This strategy removed ∼10,000 genes from both haplomes and left ∼43,000 genes in each annotation. Complete BUSCO score for this gene set decreased by ∼1% compared to the original full gene set. In the other three subsets (2-4) of genes, where less stringent criteria were applied, ∼3,000-4,000 genes were removed and complete BUSCO scores maintained above 98%. In two of the subsets where the genes with functional and gene family were taken into consideration (Subset 3 & 4), complete BUSCO scores remained the same as the original gene set even after removing thousands of genes. CROG gene count analyses were performed on the original full set, Subset 1 and Subset 3. The CROG gene count clustermaps from the three gene sets showed highly similar clustering patterns (Figure 6 and Supplemental Figure S8), indicating that removing genes did not alter the overall gene family circumscription. The average CROG gene count z-score decreased from 0.330 in the original full set, to 0.297 in Subset 3, and to 0.008 in Subset 1, indicating values closer to expectation as more rigorous evidence categories are applied.

**Table 4.**
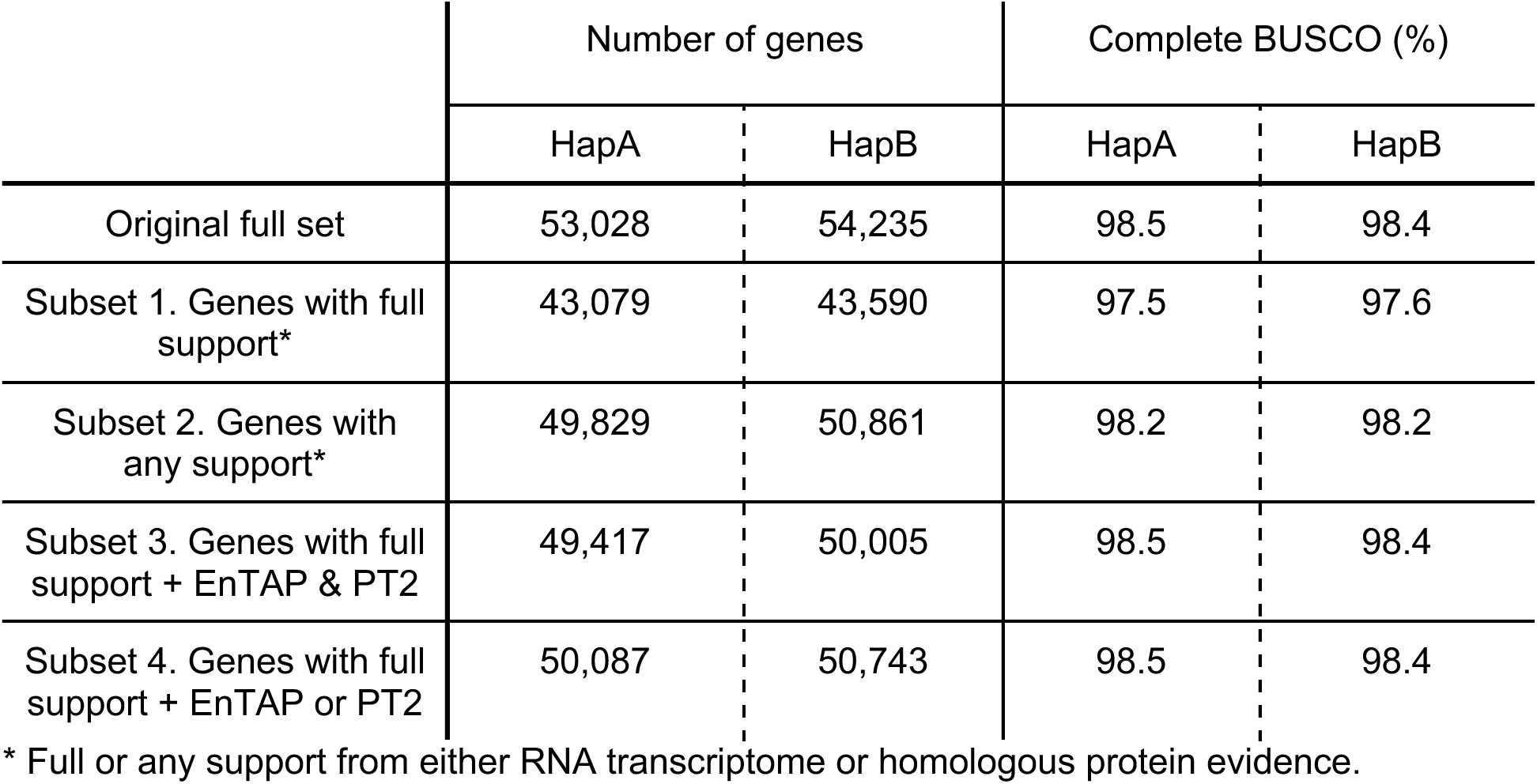
Summary of genes mapped with various evidence source and completeness assessments of those gene subsets.

### Plastid Genomes Assembly and Annotation

The chloroplast genome of the ‘WA 38’ apple is 159,915 bp in length, which is smaller than most assembled *Malus* chloroplast genomes (Naizaier *et al*. 2019; Yan *et al*. 2019; Zhao *et al*. 2019; Ha *et al*. 2020; Miao *et al*. 2022; Li *et al*. 2022a). The plastome consisted of a typical quadripartite structure with a pair of inverted repeat regions (IR) of the same length (26,352 bp) separated by a long single copy region (LSC) (88,052 bp) and a short single copy region (SSC) (19,159 bp). The IR regions and the SSC regions were all similar in length to that of other *Malus* chloroplasts (Naizaier *et al*. 2019; Yan *et al*. 2019; Zhao *et al*. 2019; Ha *et al*. 2020; Miao *et al*. 2022; Li *et al*. 2022a). A total of 134 unique genes were annotated, including 86 protein-coding genes, 42 tRNA genes, and 7 rRNA genes. Moreover, eight protein-coding genes (*ycf1, ycf2, rpl2, rpl23, ndhB, rps7, rps12, rps19-fragment*), ten tRNA genes (*trnE-UUC, trnI-GAU, trnA-UGC, trnL-CAA, trnM-CAU, trnN-GUU, trnR-ACG, trnI-CAU, trnN-GUU, trnV-GAC*), all four rRNA genes (*rrn16, rrn23, rrn4.5, rrn5*) were located wholly within the IR regions (Figure 8). Twelve protein-coding genes, eight tRNA genes, and one rRNA gene (*rrn16*) contain introns. The majority of which contained one intron (19 genes), with only two genes (*pafl* and *clpP1*) containing two introns.

**Figure 8.**
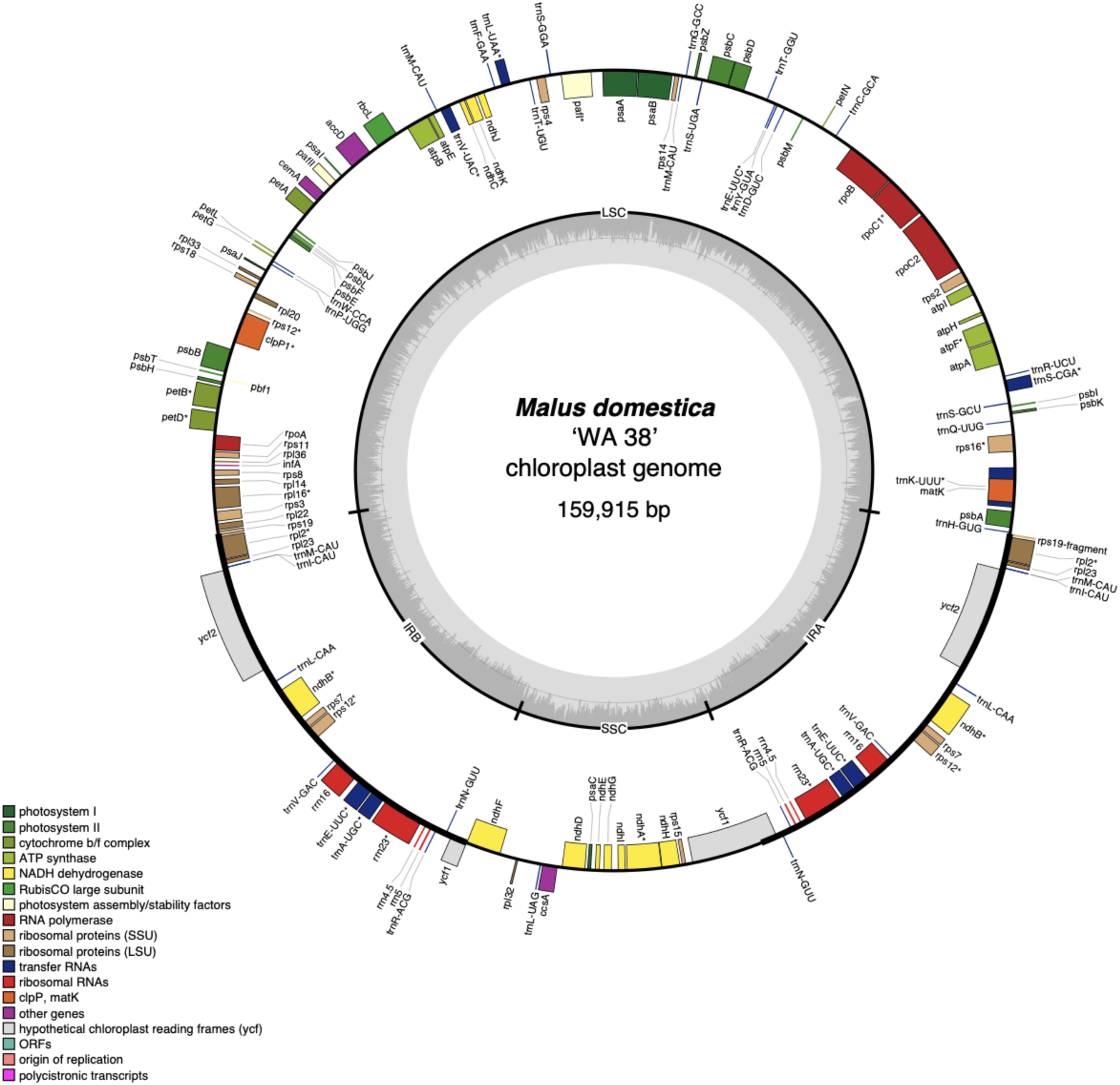
Chloroplast genome map of ‘WA 38’ with annotation. The outer circle shows the locations of genes, colored according to their function and biological pathways as shown in the figure legend. Forward-encoded genes are drawn on the outside of the circle, while reverse-encoded genes are on the inside of the circle. The middle circle shows locations of the four major sections of the chloroplast: LSC (long single copy), SSC (short single copy), IRA (inverted repeat A), and IRB (inverted repeat B). The inner gray circle shows GC content across the chloroplast genome.

The mitochondrial genome of the ‘WA 38’ apple is 451,423bp long and contains 64 annotated genes. This annotation includes 4 rRNA genes (two copies of 26S, and one copy of both 18S and 5S), 20 tRNA genes (including two copies of *trnaA-FME* and three copies of *trnaF-GAA*), and 40 protein-coding genes (including two copies of *atp1, apt8, cox3, nad6, nad7, rnaseH, rps12*, *rps3*, and *sdh4*). (Figure 9)

**Figure 9.**
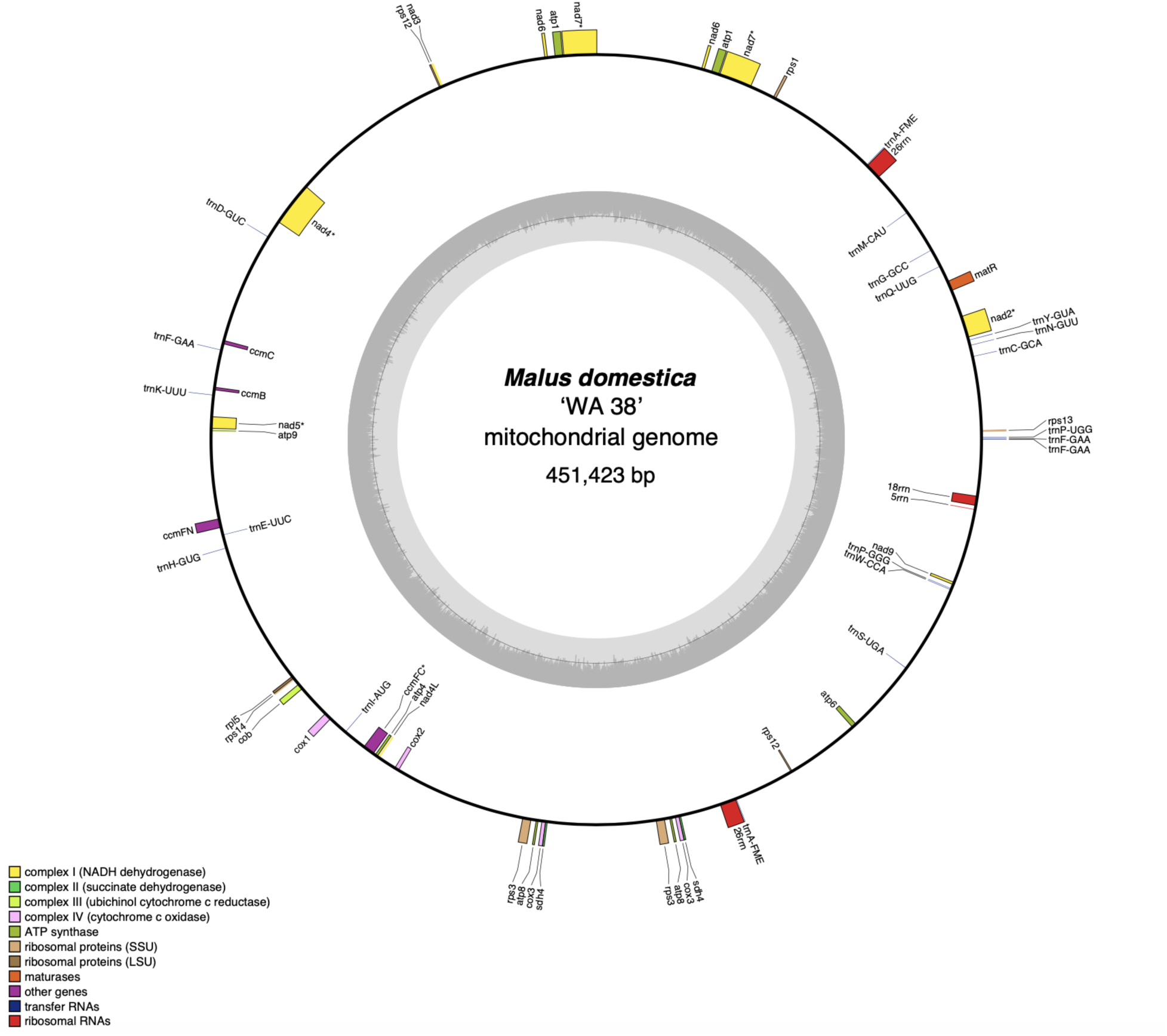
Mitochondrial genome map of ‘WA 38’ with annotation. The outer circle shows the locations of genes, colored according to their function and biological pathways as shown in the figure legend. Forward-encoded genes are drawn on the outside of the circle, while reverse-encoded genes are on the inside of the circle. The inner gray circle shows GC content across the mitochondria genome.

## Discussion

Genomes are essential resources for research communities. In order to provide accessible, hands-on training to the next generation of plant genome scientists, we engaged students in the construction of a genome for the ‘WA 38’ (Cosmic Crisp®) apple. Our guiding philosophy is ‘inclusion and novelty’, where we aim to build a high-quality reference genome that is useful to a wide range of current and future research communities.

We emphasized assembly quality by leveraging our recent ‘Honeycrisp’ genome (Khan et al. 2022) to fully resolve haplotypes, *i.e.* the specific genetic contributions of each parent are known and are represented in each respective haplome. As the first pome fruit genome to achieve this level of resolution, the ‘WA 38’ genome provides a unique resource for researchers across various fields to explore genome-scale genomic signatures that were previously unattainable for pome fruit research. Examples include a more in-depth understanding of genetic variation and inheritance, identification of alleles associated with specific traits (paving the way for allele specific expression experiments), and opportunities to perform trait association analyses with higher resolution (useful for breeding programs to identify new genetic markers linked to desirable traits) (Talbot *et al*. 2024).

We also emphasized genome annotation quality, aiming to provide a hierarchy of hypothesized gene models, where we compile a more complete list of putative genes, with increasingly stringent evidence categories allowing users to access and use the appropriate set of annotations for their application. By breaking from convention where a single stringency for genome annotation has historically been set in published genomes, our approach provides an annotation matrix that allows users to explore gene space as a function of annotation support. Our original, full gene set contains ∼54,000 putative gene models, almost 9,000 more than most other *Malus* genomes (Supplemental Table S5). Subsequent filtering using various evidence sources successfully adjusted the gene number closer to expected, although this resulted in reduced completeness in some cases (Table 4). Subset 1, where only genes with full support were selected, is the most stringent criteria we used for gene selection. Although the BUSCO completeness score dropped by ∼1%, it’s still among the highest in *Malus* annotations and the average CROG gene count z-score indicates that the overall number of genes in CROG are very close to expectation (Supplemental Figure S8). However, a collection of ‘cold’ orthogroups (containing fewer than expected number of genes compared to the rest annotations) emerged in the ‘Honeycrisp’ plus ‘WA 38’ cluster from the CROG analysis (highlighted with a box in Supplemental Figure S8). Since these cold spots were not observed in the original full gene set nor the less rigorously filtered Subset 3, and are unique to the genomes annotated with the same method and same filtering strategy, they are likely the result of a methodological bias. This subset, Subset 1, is expected to contain fewer false positives at the cost of also dropping a small amount of true positives; suitable for analysis that requires high-confidence gene models, such as reconstructing species or pedigree relationship. Subset 3, which contains all genes from Subset 1 and genes with both EnTAP and PlantTribes2 evidence, has a similar gene number to the most recently published apple genomes, namely ‘Honeycrisp’, ‘Fuji’, ‘M9’, and ‘MM106’. Subset 3 maintained the same BUSCO completeness score and did not have the ‘cold’ orthogroup observed in Subset 1. Thus, Subset 3 may contain more false positive genes, but it also retains the most true positives; suitable for most analyses that can tolerate a small amount of false positive gene models. Furthermore, similar to the ‘Honeycrisp’ plus ‘WA 38’ cluster with shared unique ‘cold’ orthogroup zones in the Subset 1 CROG analysis, genomes annotated by the same research group tend to exhibit similar gene count patterns (CROG analysis - Figure 6), suggesting that methodological bias in a seemingly subjective analysis may lead to a more similar gene landscape within those annotations. The most surprising examples are the cluster of ‘Gala’ with the two wild *Malus* progenitors (*i.e.* different species), and the cluster of *Malus fusca* with *Pyrus communis* ‘d’Anjou’ (*i.e.* different genera). In addition, although most of the published *Malus* genome annotations have a similar number of genes (∼45,000, Supplemental Table S5), the CROG analysis identified different collections of orthogroups with higher (warm color) or lower (cool color) than average gene counts across clusters. These ‘warm’ and ‘cool’ orthogroup spots are not necessarily indicative of gene family expansions or contractions (a separate analysis would be required), but does provide valuable insight into the gene space within the context of lineage-specific genome annotations and highlights potential areas for genome resource improvement. We believe the methodological bias revealed by the CROG analysis should be addressed or acknowledged before further analyses of gene family expansions and contractions in *Malus* is performed.

Throughout this project, we emphasized community engagement and enforce standardization of genome resources. The AgBioData Genome Nomenclature working group is dedicated to providing recommendations for consistent genome and gene model nomenclature that meets the FAIR data principle (Wilkinson *et al*. 2016). We worked together with this working group and the Rosaceae community genome database (Genome Database for Rosaceae, GDR, (Jung *et al*. 2019)) to improve the existing nomenclature for Rosaceae genomes. The adoption of standardized nomenclature for plant genomes represents a significant advancement in the field of plant genomics as it helps reduce confusion and potential errors, thereby enhancing the reliability and reproducibility of genomic research. In addition, we followed a previously-established gene family classification protocol (Wafula *et al*. 2022; Khan *et al*. 2022) that circumscribed genes into pre-computed orthogroups. Such a practice not only reduces computational resource requirements, but also allows researchers to more easily compare findings across studies. The uniformity, achieved by taking advantage of the already-existing community resource, facilitates clearer communication, ensuring that discoveries are accurately attributed and understood in the context of existing knowledge.

Our work emphasized the “reproducibility” of FAIR (Findable, Accessible, Interoperable and Reproducible) data. All bioinformatics analyses follow some workflow whether it is manually developed as work progresses by the researcher or is the product of an automated workflow managed by software tools like Galaxy (The Galaxy Community, 2022) (graphical interface), Nextflow (di Tommaso *et al*. 2017) or Snakemake (Mölder et al. 2021) (command-line interface). Automated workflows create reproducible analyses because the version and parameters are easily documented and software is commonly dockerized. For manually developed workflows, the process is prone to being haphazard and disorganized and difficult to share. Thus, many workflows are simply reduced to a brief description of software tools in Methods sections of journal articles with software versions and important parameters often missing. As introduced in the Results section, we provide a complete set of scripts and dockeried software to completely recreate every analysis in the assembly and annotation of the WA 38 genome. The organizational structure of the repository follows the Bioinformatics Notebook protocol developed by our team (https://gitlab.com/ficklinlab-public/bioinformatics-notebook/). The goal of this protocol is to ensure that complex manually executed workflows can be shared for reproducibility, the format is readable by others and backups of critical data are supported.

Briefly, the directories are ordered using a numeric prefix indicating the order that analyses should be performed. Inside each directory are sub-directories with smaller tasks. For each task all relevant scripts and instructions are provided. All software used by the project is dockerized and scripts contain the full parameter set used for every step. While there are areas for improvement, the protocol, when followed, allows for easy sharing of the workflow via a Git repository. In our view, this approach is a novel contribution towards FAIR data by ensuring that non-automated workflows can be shared and are fully reproducible.

In addition to providing a fully reproducible workflow for the assembly of the ‘WA 38’ genome. We generalized the scripts for any genome assembly and shared those as part of the three ACTG GtiHub repositories mentioned in the Results setion. The new ACTG general workflow is designed to provide training that is applicable for a wide range of species. The ACTG repositories are a work in progress as we seek to create a generic, species-agnostic workflow that will serve the broader American Campus Tree Genome (ACTG) community.

## Availability of source code and requirements

Project name: ‘WA 38’ whole genome assembly and annotation

Project home page: https://gitlab.com/ficklinlab-public/wa-38-genome

Operating system(s): Platform independent

Programming language: bash, python, awk, perl

Other requirements: singularity, nextflow, java, python

License: Not applicable

Any restrictions to use by non-academics: No restrictions

RRID: Not applicable

## Data Availability

Raw reads generated for this project are publicly available at NCBI under BioProject: PRJNA1072127. Genome assembly and annotation are available on GDR: https://www.rosaceae.org/Analysis/20220983

## Authors’ contributions

L.H and S.F acquired funding for this project. All authors contributed to data analysis, data interpretation, and manuscript writing.

## References

Andrews. S, 2010 FastQC: A Quality Control Tool for High Throughput Sequence Data [Online]. Available online at: http://www.bioinformatics.babraham.ac.uk/projects/fastqc/

NC State Extension, n.d. Apples - Malus domestica, North Carolina Extension Gardener Plant Toolbox. Retrieved May 23, 2024 [online] https://plants.ces.ncsu.edu/plants/malus-domestica/common-name/apples/

Brůna, T., K. J. Hoff, A. Lomsadze, M. Stanke, and M. Borodovsky, 2021 BRAKER2: automatic eukaryotic genome annotation with GeneMark-EP+ and AUGUSTUS supported by a protein database. NAR Genom Bioinform 3: lqaa108.

Chen C., Wu Y., Li J., Wang X., Zeng Z., Xu J., Liu Y., Feng J., Chen H., He Y., and Xia R., 2023 TBtools-II: A “one for all, all for one” bioinformatics platform for biological big-data mining. Mol. Plant 16: 1733–1742.

Cheng, H., G. T. Concepcion, X. Feng, H. Zhang, and H. Li, 2021 Haplotype-resolved de novo assembly using phased assembly graphs with hifiasm. Nat. Methods 18: 170–175.

Chen, X., S. Li, D. Zhang, M. Han, X. Jin et al., 2019 Sequencing of a Wild Apple (*Malus baccata*) Genome Unravels the Differences Between Cultivated and Wild Apple Species Regarding Disease Resistance and Cold Tolerance. G3 9: 2051–2060.

Chen, S., Y. Zhou, Y. Chen, and J. Gu, 2018 fastp: an ultra-fast all-in-one FASTQ preprocessor. Bioinformatics 34: i884–i890.

Choi, J. Y., L. R. Abdulkina, J. Yin, I. B. Chastukhina, J. T. Lovell et al., 2021 Natural variation in plant telomere length is associated with flowering time. Plant Cell 33: 1118–1134.

Brown, M., P. M. González De la Rosa, and M. Blaxter, (2023). A Telomere Identification Toolkit (v0.2.41). Zenodo. 10.5281/zenodo.10091385

Crosby, J. A., J. Janick, P. C. Pecknold, J. C. Goffreda, and S. S. Korban, 1994 ‘Enterprise’ Apple. HortScience 29: 825–826.

Daccord, N., J.-M. Celton, G. Linsmith, C. Becker, N. Choisne et al., 2017 High-quality de novo assembly of the apple genome and methylome dynamics of early fruit development. Nat. Genet. 49: 1099–1106.

Danecek, P., J. K. Bonfield, J. Liddle, J. Marshall, V. Ohan et al., 2021 Twelve years of SAMtools and BCFtools. Gigascience 10: giab008.

di Tommaso, P., M. Chatzou, E. W. Floden, P. P. Barja, E. Palumbo, and C. Notredame, 2017. Nextflow enables reproducible computational workflows. Nature Biotechnology, 35(4), 316–319.

Dierckxsens, N., P. Mardulyn, and G. Smits, 2017 NOVOPlasty: de novo assembly of organelle genomes from whole genome data. Nucleic Acids Res. 45: e18.

Di Guardo, M., A. Tadiello, B. Farneti, G. Lorenz, D. Masuero et al., 2013 A multidisciplinary approach providing new insight into fruit flesh browning physiology in apple (*Malus* x *domestica* Borkh.). PLoS One 8: e78004.

Dong, X., Z. Wang, L. Tian, Y. Zhang, D. Qi et al., 2020 De novo assembly of a wild pear (*Pyrus betuleafolia*) genome. Plant Biotechnol. J. 18: 581–595.

Doyle, J. J., and J. L. Doyle, 1987 A rapid DNA isolation procedure for small quantities of fresh leaf tissue: RESEARCH.

Durand, N. C., M. S. Shamim, I. Machol, S. S. P. Rao, M. H. Huntley et al., 2016 Juicer Provides a One-Click System for Analyzing Loop-Resolution Hi-C Experiments. Cell Syst 3: 95–98.

Earl, D., K. Bradnam, J. St John, A. Darling, D. Lin et al., 2011 Assemblathon 1: a competitive assessment of de novo short read assembly methods. Genome Res. 21: 2224–2241.

Evans, K. M., B. H. Barritt, B. S. Konishi, L. J. Brutcher, and C. F. Ross, 2012 “WA 38” Apple. HortScience 47: 1177–1179.

FGN, 2020 Global apple market reached $78B: set to continue moderate growth. Fruit Growers News. Retrieved May 23, 2024 [online] https://fruitgrowersnews.com/news/global-apple-market-reached-78m-set-to-continue-moderate-growth/

Gabriel, L., K. J. Hoff, T. Brůna, M. Borodovsky, and M. Stanke, 2021 TSEBRA: transcript selector for BRAKER. BMC Bioinformatics 22: 566.

Goremykin, V. V., P. J. Lockhart, R. Viola, and R. Velasco, 2012 The mitochondrial genome of *Malus domestica* and the import-driven hypothesis of mitochondrial genome expansion in seed plants. Plant J. 71: 615–626.

Grabherr, M. G., B. J. Haas, M. Yassour, J. Z. Levin, D. A. Thompson et al., 2011 Full-length transcriptome assembly from RNA-Seq data without a reference genome. Nat. Biotechnol. 29: 644–652.

Greiner, S., P. Lehwark, and R. Bock, 2019 OrganellarGenomeDRAW (OGDRAW) version 1.3.1: expanded toolkit for the graphical visualization of organellar genomes. Nucleic Acids Res. 47: W59–W64.

Haas, B. J., A. L. Delcher, S. M. Mount, J. R. Wortman, R. K. Smith Jr et al., 2003 Improving the Arabidopsis genome annotation using maximal transcript alignment assemblies. Nucleic Acids Res. 31: 5654–5666.

Hadish, J. A., T. D. Biggs, B. T. Shealy, M. R. Bender, C. B. McKnight et al., 2022 GEMmaker: process massive RNA-seq datasets on heterogeneous computational infrastructure. BMC Bioinformatics 23: 1–11.

Ha, Y.-H., B. Maisupova, K. Choi, H.-J. Kim, D. Dosmanvetov et al., 2020 Report on a complete chloroplast genome sequence of wild apple tree, *Malus sieversii* (Lebed.) M. Roem. Mitochondrial DNA B Resour. 5: 1504–1505.

Harkess, A., 2022 The American Campus Tree Genomes Documentation. Retrieved May 23, 2024 [online] https://actg-wgaa.readthedocs.io/en/latest/index.html

Harris, R.S., 2007 Improved pairwise alignment of genomic DNA. Ph.D. Thesis, The Pennsylvania State University. Retrieved May 23, 2024 [online] https://www.bx.psu.edu/~rsharris/rsharris_phd_thesis_2007.pdf

Hart, A. J., S. Ginzburg, M. (sam) Xu, C. R. Fisher, N. Rahmatpour et al., 2020 EnTAP: Bringing faster and smarter functional annotation to non-model eukaryotic transcriptomes. Mol. Ecol. Resour. 20: 591–604.

Hoff, K. J., S. Lange, A. Lomsadze, M. Borodovsky, and M. Stanke, 2016 BRAKER1: Unsupervised RNA-Seq-Based Genome Annotation with GeneMark-ET and AUGUSTUS. Bioinformatics 32: 767–769.

Howard, N. P., E. van de Weg, D. S. Bedford, C. P. Peace, S. Vanderzande et al., 2017 Elucidation of the ‘Honeycrisp’ pedigree through haplotype analysis with a multi-family integrated SNP linkage map and a large apple (*Malus×domestica*) pedigree-connected SNP data set. Horticulture Research 4: 17003.

Huerta-Cepas, J., D. Szklarczyk, D. Heller, A. Hernández-Plaza, S. K. Forslund et al., 2019 eggNOG 5.0: a hierarchical, functionally and phylogenetically annotated orthology resource based on 5090 organisms and 2502 viruses. Nucleic Acids Res. 47: D309–D314.

Johnston, J. W., and P. Brookfield, 2012 Delivering postharvest handling protocols for apples and pears faster: Integrating “omics” and physiology approaches. Acta Hortic. 23–28.

Jung, S., T. Lee, C.-H. Cheng, K. Buble, P. Zheng et al., 2019 15 years of GDR: New data and functionality in the Genome Database for Rosaceae. Nucleic Acids Res. 47: D1137–D1145.

Khan, A., S. B. Carey, A. Serrano, H. Zhang, H. Hargarten et al., 2022 A phased, chromosome-scale genome of “Honeycrisp” apple (*Malus domestica*). GigaByte 2022: gigabyte69.

Kurtz, S., A. Phillippy, A. L. Delcher, M. Smoot, M. Shumway et al., 2004 Versatile and open software for comparing large genomes. Genome Biol. 5: R12.

Kuznetsov, D., F. Tegenfeldt, M. Manni, M. Seppey, M. Berkeley et al., 2022 OrthoDB v11: annotation of orthologs in the widest sampling of organismal diversity. Nucleic Acids Res. 51: D445–D451.

Li, X., Z. Ding, H. Miao, J. Bao, and X. Tian, 2022a Complete chloroplast genome studies of different apple varieties indicated the origin of modern cultivated apples from and. PeerJ 10: e13107.

Li, H., and R. Durbin, 2009 Fast and accurate short read alignment with Burrows-Wheeler transform. Bioinformatics 25: 1754–1760.

Liebhard, R., M. Kellerhals, W. Pfammatter, M. Jertmini, and C. Gessler, 2003 Mapping quantitative physiological traits in apple (*Malus* x *domestica* Borkh.). Plant Mol. Biol. 52: 511–526.

Li, X., L. Kui, J. Zhang, Y. Xie, L. Wang et al., 2016 Improved hybrid de novo genome assembly of domesticated apple (*Malus* x *domestica*). Gigascience 5: 35.

Li, W., C. Chu, H. Li, et al., 2024 Near-gapless and haplotype-resolved apple genomes provide insights into the genetic basis of rootstock-induced dwarfing. Nat Genet 56, 505–516.

Li, W., J. Liu, H. Zhang, Z. Liu, Y. Wang et al., 2022b Plant pan-genomics: recent advances, new challenges, and roads ahead. J. Genet. Genomics 49: 833–846.

Li, Y., M. Pi, Q. Gao, Z. Liu, and C. Kang, 2019 Updated annotation of the wild strawberry *Fragaria vesca* V4 genome. Horticulture Research 6: 1–9.

Lovell, J. T., A. Sreedasyam, M. Eric Schranz, M. Wilson, J. W. Carlson et al., 2022 GENESPACE tracks regions of interest and gene copy number variation across multiple genomes.

Lum, G. B., B. J. Shelp, J. R. DeEll, and G. G. Bozzo, 2016 Oxidative metabolism is associated with physiological disorders in fruits stored under multiple environmental stresses. Plant Sci. 245: 143–152.

Manni, M., M. R. Berkeley, M. Seppey, F. A. Simão, and E. M. Zdobnov, 2021 BUSCO Update: Novel and Streamlined Workflows along with Broader and Deeper Phylogenetic Coverage for Scoring of Eukaryotic, Prokaryotic, and Viral Genomes. Mol. Biol. Evol. 38: 4647–4654.

Mansfeld, B. N., A. Yocca, S. Ou, A. Harkess, E. Burchard et al., 2023 A haplotype resolved chromosome-scale assembly of North American wild apple *Malus fusca* and comparative genomics of the fire blight Mfu10 locus. Plant J. 116: 989–1002.

Marçais, G., and C. Kingsford, 2011 A fast, lock-free approach for efficient parallel counting of occurrences of k-mers. Bioinformatics 27: 764–770.

Mendoza, M., Hanrahan, I., & Bolaños, G., 2022 2022 Update: Additional WA 38 harvest and storage considerations. Retrieved May 23, 2024 [online] https://treefruit.wsu.edu/article/2022-update-additional-wa-38-harvest-and-storage-consideration

Miao, H., J. Bao, X. Li, Z. Ding, and X. Tian, 2022 Comparative analyses of chloroplast genomes in “Red Fuji” apples: low rate of chloroplast genome mutations. PeerJ 10: e12927.

Mölder, F., K. P. Jablonski, B. Letcher, M. B. Hall, C. H. Tomkins-Tinch et al. 2021 Sustainable data analysis with Snakemake [version 2; peer review: 2 approved]. F1000Research, 10:33

Naizaier, R., Z. Qu, S. Wu, and X. Tian, 2019 The complete chloroplast genome of Malus *sieversii* (Rosaceae), a wild apple tree in Xinjiang, China. Mitochondrial DNA B Resour. 4: 983–984.

Nattestad, M., and M. C. Schatz, 2016 Assemblytics: a web analytics tool for the detection of variants from an assembly. Bioinformatics 32: 3021–3023.

NCBI Organelle genome resources, n.d. Retrieved May 23, 2024 [online] https://www.ncbi.nlm.nih.gov/genome/organelle/

O’Leary, N. A., M. W. Wright, J. R. Brister, S. Ciufo, D. Haddad et al., 2016 Reference sequence (RefSeq) database at NCBI: current status, taxonomic expansion, and functional annotation. Nucleic Acids Res. 44: D733–45.

Ou, S., W. Su, Y. Liao, K. Chougule, J. R. A. Agda et al., 2019 Benchmarking transposable element annotation methods for creation of a streamlined, comprehensive pipeline. Genome Biol. 20: 1–18.

Pareek, S. T. de F. S., 2019 Postharvest physiological disorders in fruits and vegetables (S. Tonetto de Freitas & S. Pareek, Eds.). CRC Press, Boca Raton : Taylor & Francis, 2018. phasegenomics, n.d. Hic_QC: a (very) simple script to QC Hi-C data. GitHub. https://github.com/phasegenomics/hic_qc

Quinlan, A. R., and I. M. Hall, 2010 BEDTools: a flexible suite of utilities for comparing genomic features. Bioinformatics 26: 841–842.

Raymond, O., J. Gouzy, J. Just, H. Badouin, M. Verdenaud et al., 2018 The *Rosa* genome provides new insights into the domestication of modern roses. Nat. Genet. 50: 772–777.

Rhie, A., Walenz, B.P., Koren, S. et al., 2020 Merqury: reference-free quality, completeness, and phasing assessment for genome assemblies. Genome Biol 21, 245.

Robinson, J. T., H. Thorvaldsdóttir, W. Winckler, M. Guttman, E. S. Lander et al., 2011 Integrative genomics viewer. Nat. Biotechnol. 29: 24–26.

Robinson, J. T., D. Turner, N. C. Durand, H. Thorvaldsdóttir, J. P. Mesirov et al., 2018 Juicebox.js Provides a Cloud-Based Visualization System for Hi-C Data. Cell Syst 6: 256–258.e1.

Sallato, B., M. D. Whiting, and J. Munguia, 2021 Rootstock and nutrient imbalance leads to “Green Spot” development in ‘WA 38’ apples. HortScience 56: 1542–1548.

Serra, S., Goke, A., Sheick, R., Mendoza, M., Schmidt, T., Hanrahan, I., Ross, C., & Musacchi, S., 2023 Effects of harvest timing on maturity, fruit quality, and consumer acceptance of ‘WA 38’ apples. Acta Horticulturae, 1366, 61–68.

Sharman, S., n.d. Learning from the trees: American Campus Tree Genomes Project pushes equality in genomic science. Retrieved May 23, 2024 [online] https://hudsonalpha.shorthandstories.com/learning-from-the-trees/

Sheick, R., S. Serra, S. Musacchi, and D. Rudell, 2023 Metabolic fingerprint of ‘WA 38’ green spot symptoms reveals increased production of epicuticular metabolites by parenchyma. Sci. Hortic. 321: 112257.

Sheick, R., S. Serra, D. Rudell, and S. Musacchi, 2022 Investigations of multiple approaches to reduce Green Spot incidence in ‘WA 38’ apple. Agronomy (Basel) 12: 2822.

Shirasawa, K., A. Itai, and S. Isobe, 2021 Chromosome-scale genome assembly of Japanese pear (Pyrus pyrifolia) variety ‘Nijisseiki’ DNA Res. 28:dsab001

Smit, AFA, Hubley, R & Green, P., n.d. RepeatMasker Open-4.0.2013-2015 Retrieved May 23, 2024 [online] http://www.repeatmasker.org

Sun, X., C. Jiao, H. Schwaninger, C. T. Chao, Y. Ma et al., 2020 Phased diploid genome assemblies and pan-genomes provide insights into the genetic history of apple domestication. Nat. Genet. 52: 1423–1432.

Su, Y., X. Yang, Y. Wang, J. Li, Q. Long et al., 2024 Phased Telomere-to-Telomere Reference Genome and Pangenome Reveal an Expansion of Resistance Genes during Apple Domestication. Plant Physiol. kiae258

Talbot, S. C., K. J. Vining, J. W. Snelling, J. Clevenger, and S. A. Mehlenbacher, 2024 A haplotype-resolved chromosome-level assembly and annotation of European hazelnut (C. avellana cv. Jefferson) provides insight into mechanisms of eastern filbert blight resistance. G3. jkae021

The Galaxy Community, The Galaxy platform for accessible, reproducible and collaborative biomedical analyses: 2022 update. Nucleic Acids Research. 50: W345–W351.

Tillich, M., P. Lehwark, T. Pellizzer, E. S. Ulbricht-Jones, A. Fischer et al., 2017 GeSeq - versatile and accurate annotation of organelle genomes. Nucleic Acids Res. 45: W6–W11. trinityrnaseq, n.d. get_longest_isoform_seq_per_trinity_gene.pl. GitHub. Retrieved May 23, 2024 [online] https://github.com/trinityrnaseq/trinityrnaseq/blob/master/util/misc/get_longest_isoform_seq_per_trinity_gene.pl

Truscott, S., 2023 WSU’s Cosmic Crisp® joins top 10 bestselling U.S. apple varieties. CAHNRS News Washington State University. Retrieved May 23, 2024 [online] https://news.cahnrs.wsu.edu/article/wsus-cosmic-crisp-joins-top-10-bestselling-u-s-apple-varieties/

Uliano-Silva, M., J. G. R. N. Ferreira, K. Krasheninnikova, G. Formenti, L. Abueg et al., 2023 MitoHiFi: a python pipeline for mitochondrial genome assembly from PacBio high fidelity reads. BMC Bioinformatics 24: 1–13.

USApple, 2024 Industry at a glance Retrieved May 23, 2024 [online], https://usapple.org/industry-at-a-glance

VanBuren, R., C. M. Wai, M. Colle, J. Wang, S. Sullivan et al., 2018 A near complete, chromosome-scale assembly of the black raspberry (*Rubus occidentalis*) genome. Gigascience 7.:

Velasco, R., A. Zharkikh, J. Affourtit, A. Dhingra, A. Cestaro et al., 2010 The genome of the domesticated apple (*Malus* × *domestica* Borkh.). Nat. Genet. 42: 833–839.

Verde, I., J. Jenkins, L. Dondini, S. Micali, G. Pagliarani et al., 2017 The Peach v2.0 release: high-resolution linkage mapping and deep resequencing improve chromosome-scale assembly and contiguity. BMC Genomics 18: 225.

Vurture, G. W., F. J. Sedlazeck, M. Nattestad, C. J. Underwood, H. Fang et al., 2017 GenomeScope: fast reference-free genome profiling from short reads. Bioinformatics 33: 2202–2204.

Wafula, E. K., H. Zhang, G. Von Kuster, J. H. Leebens-Mack, L. A. Honaas et al., 2022 PlantTribes2: Tools for comparative gene family analysis in plant genomics. Front. Plant Sci. 13: 1011199.

Washington Apple Commission, 2021 Did you know? Apple Facts. Retrieved May 23, 2024 [online] https://waapple.org/did-you-know/

Wilkinson, M. D., M. Dumontier, I. J. J. Aalbersberg, G. Appleton, M. Axton et al., 2016 The FAIR Guiding Principles for scientific data management and stewardship. Sci Data 3: 160018.

Wood, D. E., and S. L. Salzberg., 2014 Kraken: ultrafast metagenomic sequence classification using exact alignments. Genome Biol 15, R46

Yan, M., X. Zhao, J. Zhou, Y. Huo, Y. Ding et al., 2019 The complete chloroplast genome of cultivated apple (*Malus domestica* Cv. “Yantai Fuji 8”). Mitochondrial DNA B Resour. 4: 1213–1216.

Yocca, A., M. Akinyuwa, N. Bailey, B. Cliver, H. Estes et al., 2024 A chromosome-scale assembly for ‘d’Anjou’ pear. G3 14.3, jkae003

Zhang, H., E. K. Wafula, J. Eilers, A. E. Harkess, P. E. Ralph et al., 2022 Building a foundation for gene family analysis in Rosaceae genomes with a novel workflow: A case study in Pyrus architecture genes. Front. Plant Sci. 13: 975942.

Zhao, X., M. Yan, Y. Ding, X. Chen, and Z. Yuan, 2019 The complete chloroplast genome of apple rootstock ‘M9.’ Mitochondrial DNA B Resour 4: 2187–2188.

Zhou, C., S. A. McCarthy, and R. Durbin, 2022 YaHS: yet another Hi-C scaffolding tool. Bioinformatics 39: btac808.

