## Supplemental Figure S1-8 for "A Haplotype-resolved, Chromosome-scale Genome for *Malus domestica* Borkh. *‘*WA 38’"

Supplemental Figure 1. Workflow of the project and each step, including A) summary workflow, B) quality control workflow, C) nuclear assembly workflow, D) Structural annotation workflow, E) Function Annotation workflow, F) Comparative Analysis Workflow, G) Chloroplast Assembly & Annotation Workflow

**A) Summary Workflow**

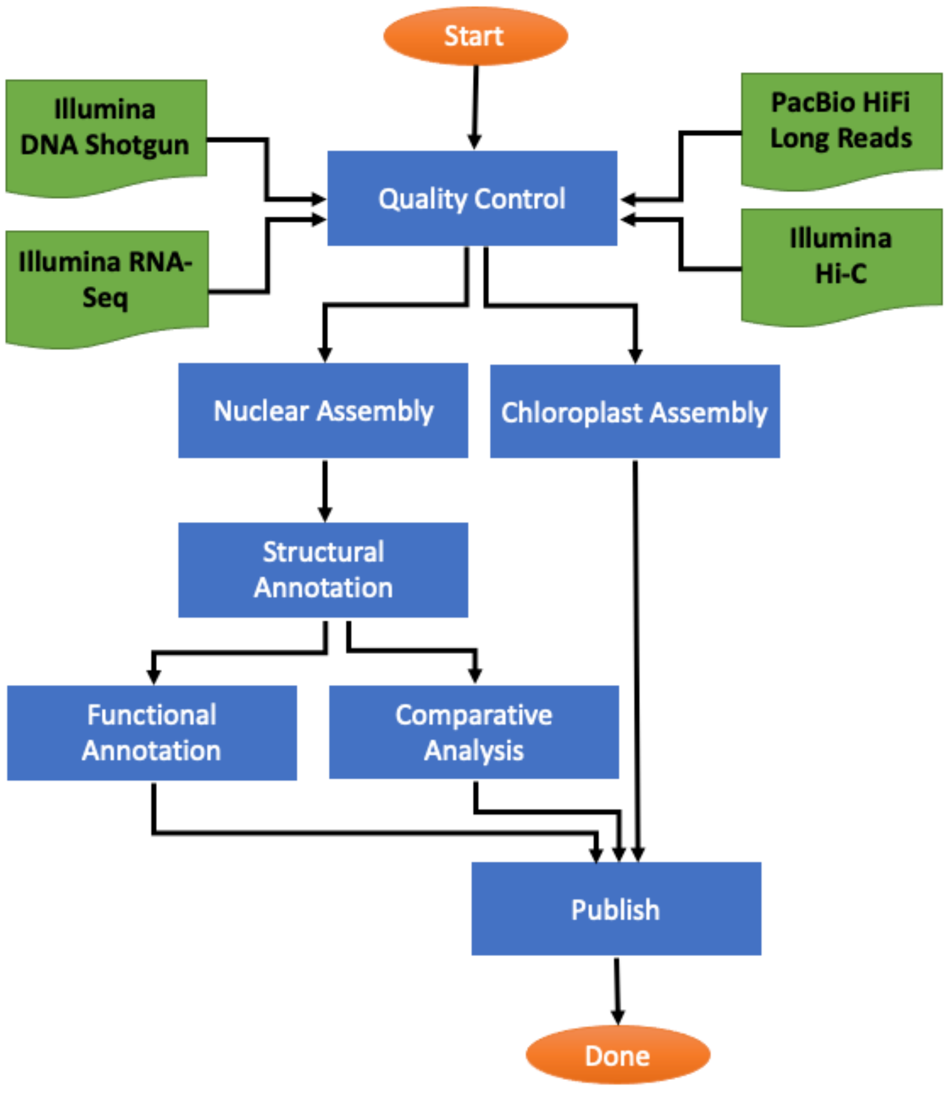

### B) Quality Control Workflow

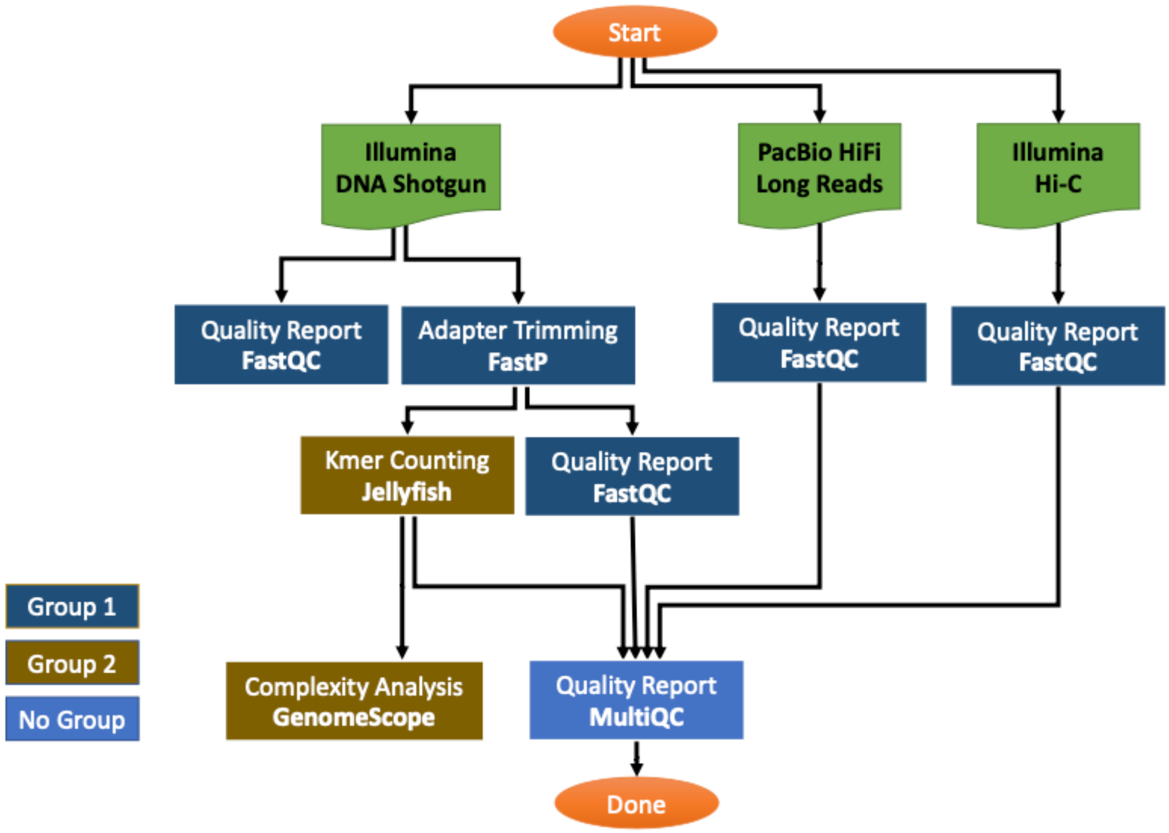

C) Nuclear Assembly Workflow

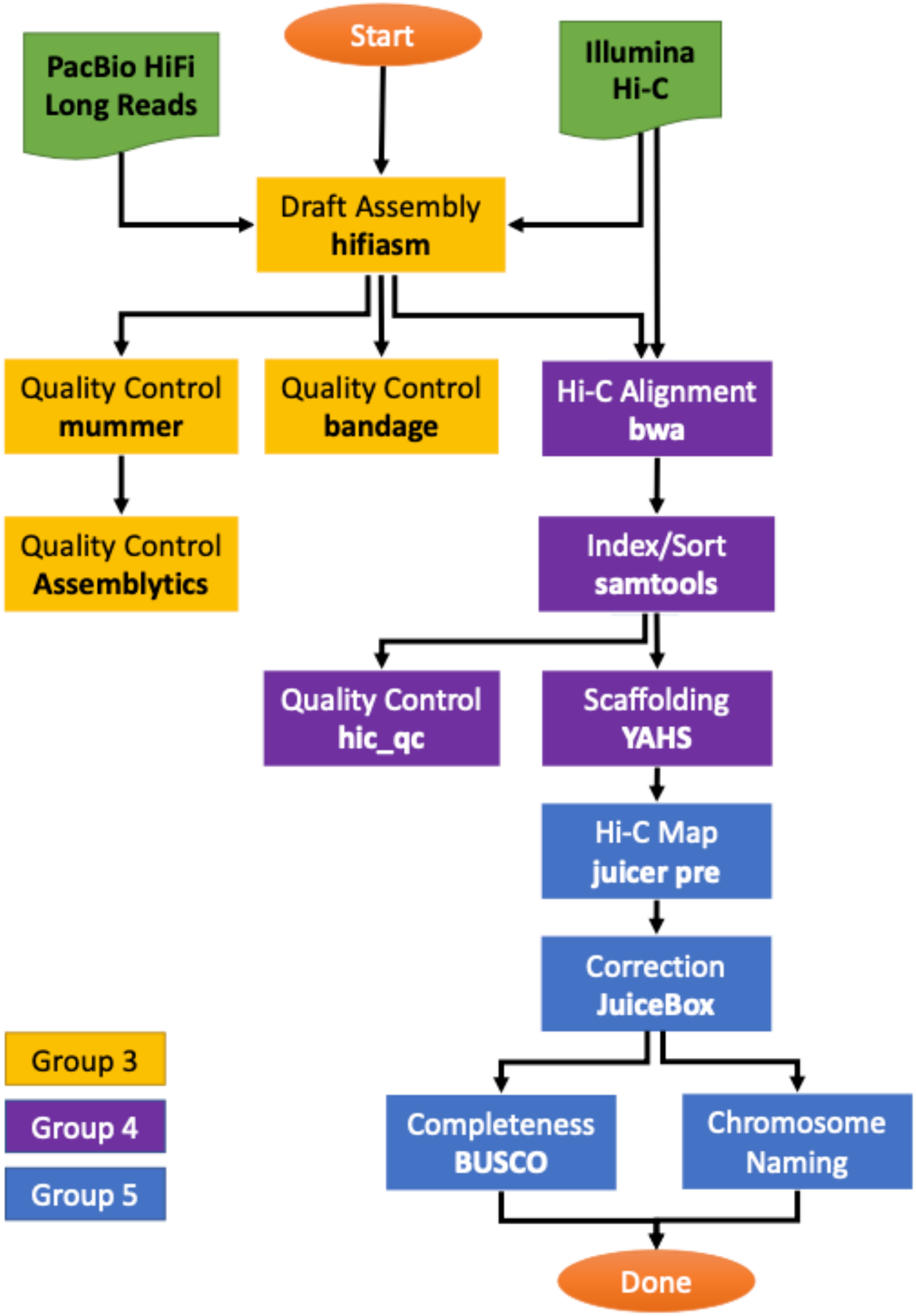

### D) Structural Annotation workflow

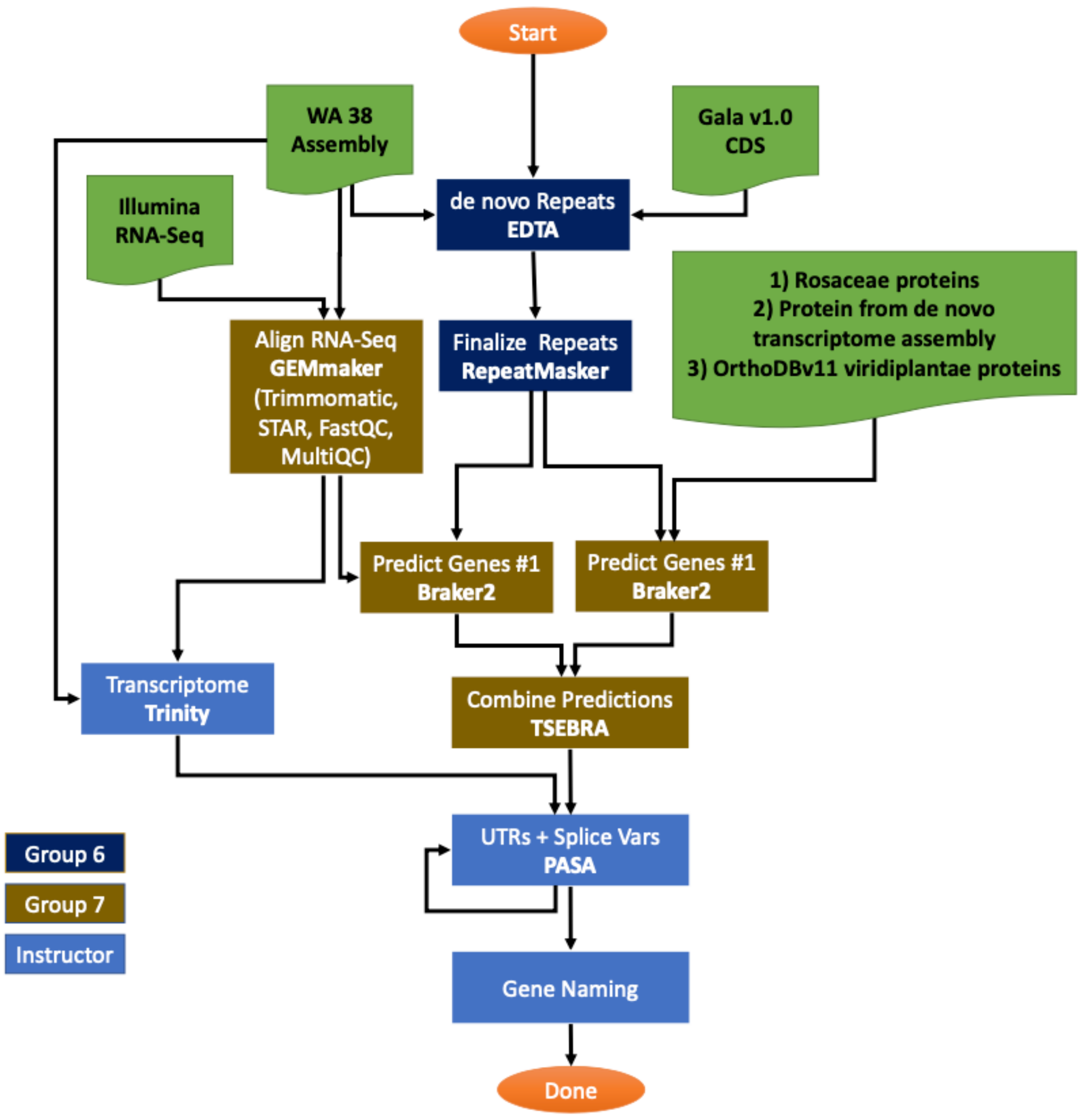

E) Function Annotation workflow

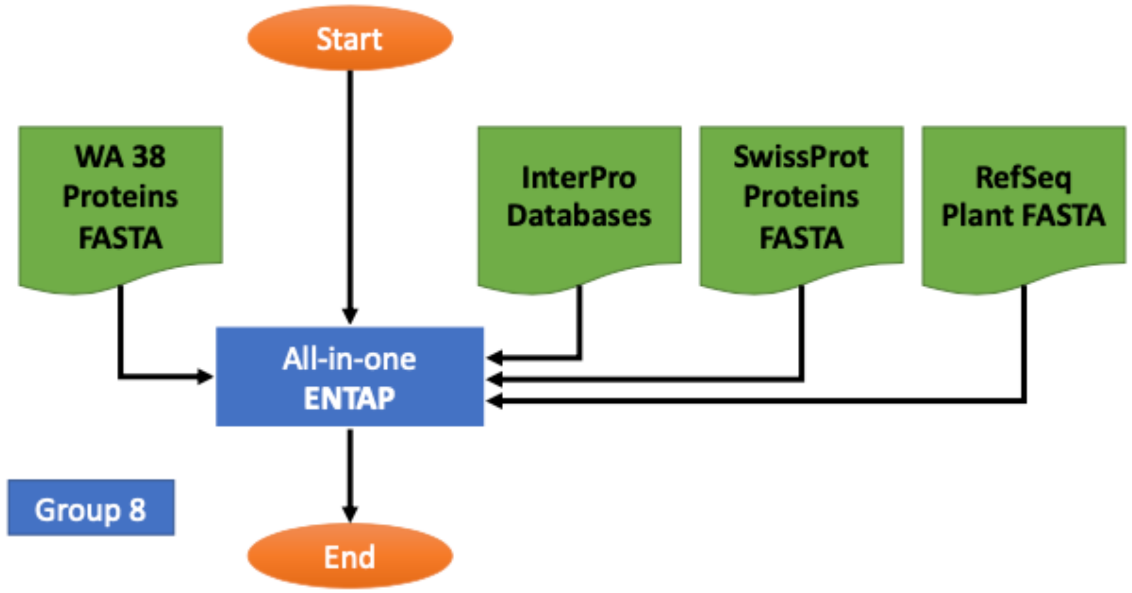


F) Comparative Analysis Workflow

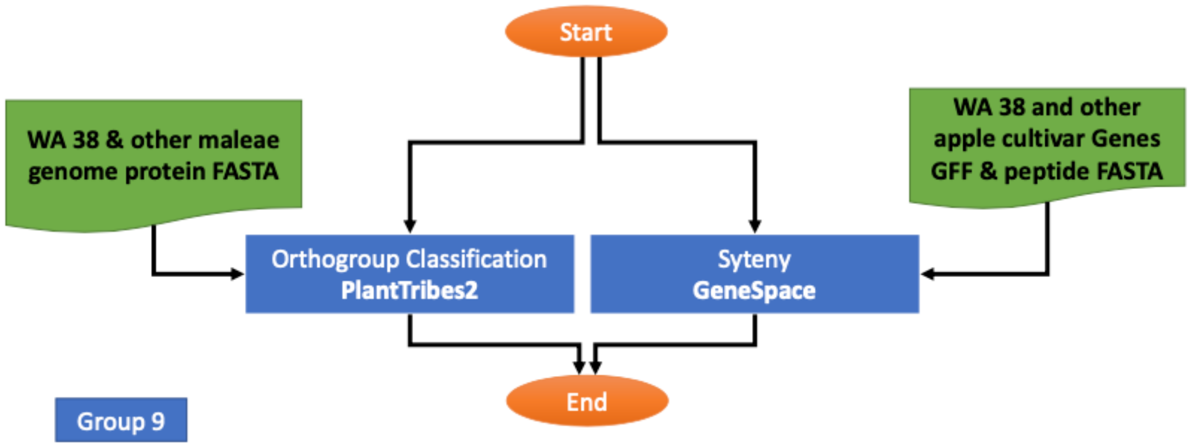

**G) Chloroplast Assembly & Annotation Workflow**

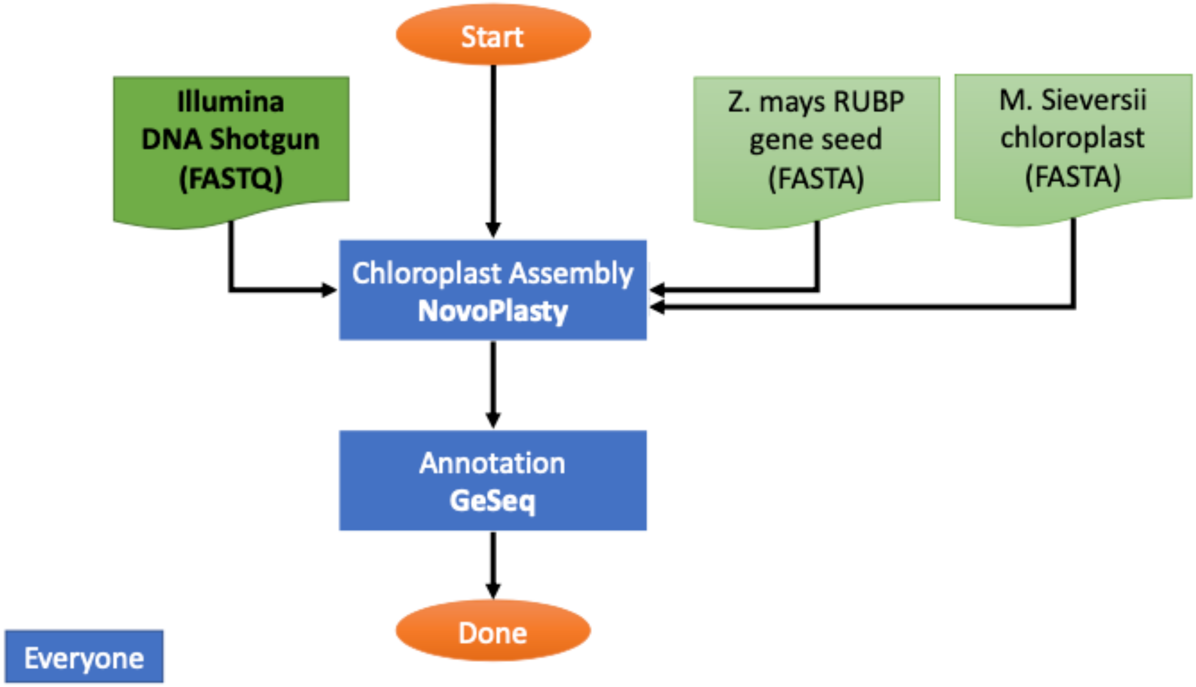

Supplemental Figure S6. Chromosome re-orientation.
LASTZ alignment graph of original ‘WA 38’ chromosomes (y axis) against corresponding ‘Gala’ chromosome (x axis). Chr#A/B_cultivar indicates the sequence in the orientation, Chr#A/B_reoriented_cultivar indicates the chromosome is reoriented according to the corresponding ‘Gala’ chromosome orientation. Blue lines indicate sequences matching in the same orientation, while red lines indicates sequences matching in reverse orientation. Only straight continuous diagonal lines indicate alignment of the entire chromosome.

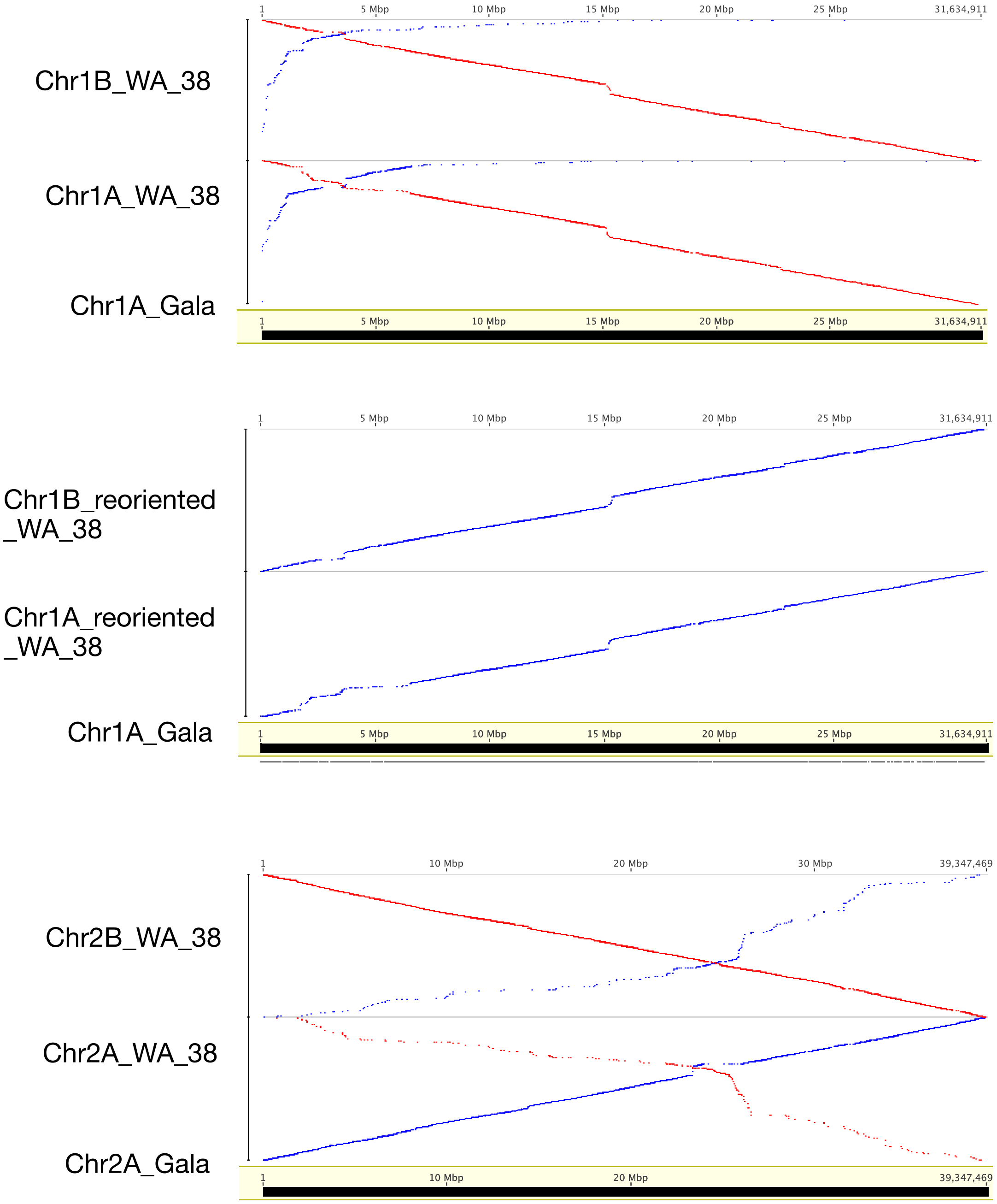

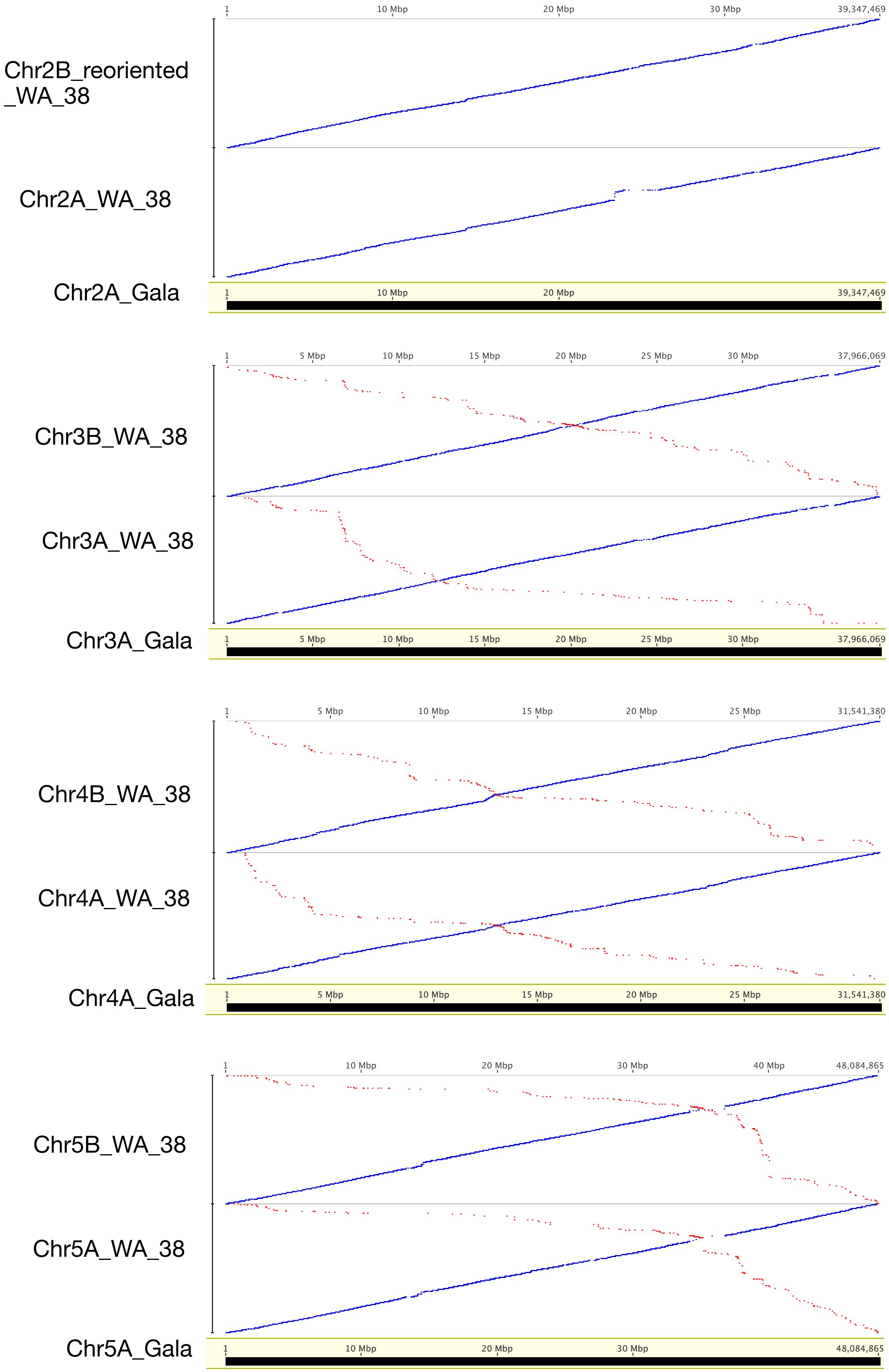

Chr2B_reoriented _WA_38
Chr2A_WA_38
Chr2A_Gala
1
10 Mbp
20 Mbp
30 Mbp
39,347,469
Chr3B_WA_38
Chr3A_WA_38
Chr3A_Gala
5 Mbp
10 Mbp
15 Mbp
20 Mbp
25 Mbp
30 Mbp
37,966,069
Chr4B_WA_38
Chr4A_WA_38
Chr4A_Gala
5 Mbp
10 Mbp
15 Mbp
20 Mbp
25 Mbp
31,541,380
Chr5B_WA_38
Chr5A_WA_38
Chr5A_Gala
10 Mbp
20 Mbp
30 Mbp
40 Mbp
48,084,865

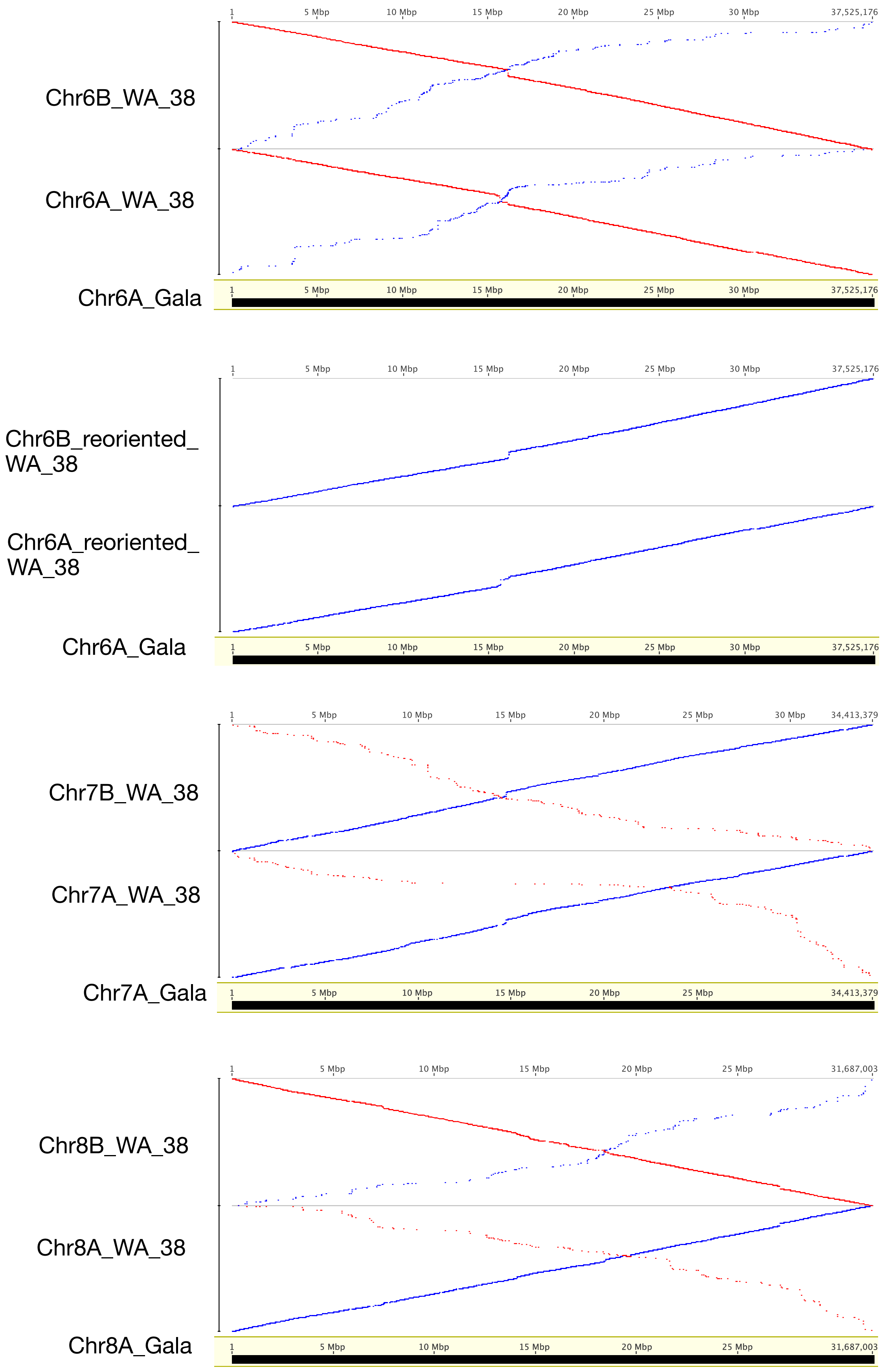

Chr6B_WA_38
Chr6A_WA_38
Chr6A_Gala
1
5 Mbp
10 Mbp
15 Mbp
20 Mbp
25 Mbp
30 Mbp
37,525,176
Chr6B_reoriented_WA_38
Chr6A_reoriented_WA_38
Chr6A_Gala
Chr7B_WA_38
Chr7A_WA_38
Chr7A_Gala
34,413,379
Chr8B_WA_38
Chr8A_WA_38
Chr8A_Gala
31,687,003

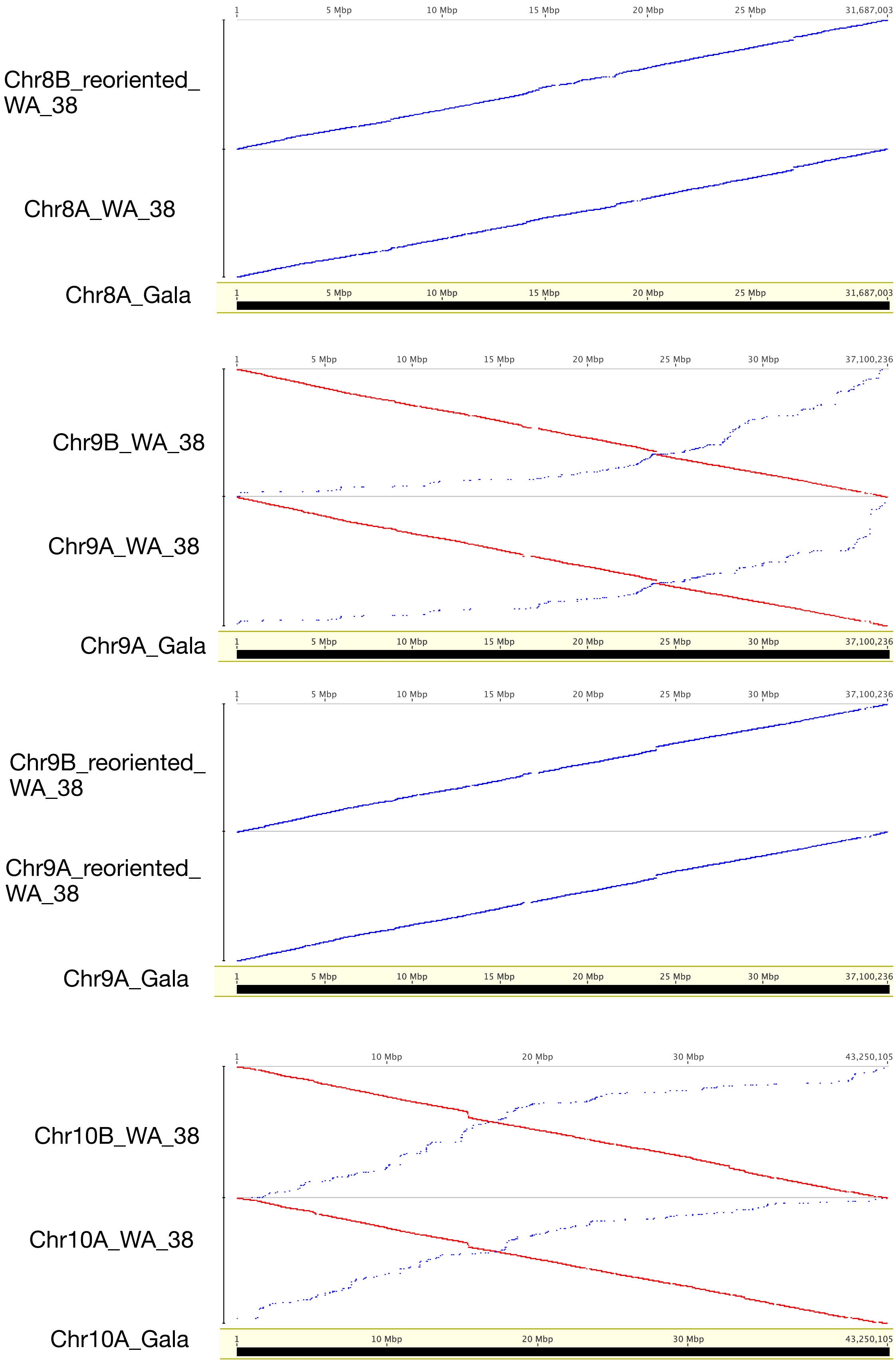

Chr8B_reoriented_WA_38
Chr8A_WA_38
Chr8A_Gala
1
5 Mbp
10 Mbp
15 Mbp
20 Mbp
25 Mbp
31,687,003
Chr9B_WA_38
Chr9A_WA_38
Chr9A_Gala
30 Mbp
37,100,236
Chr9B_reoriented_WA_38
Chr9A_reoriented_WA_38
Chr9A_Gala
Chr10B_WA_38
Chr10A_WA_38
Chr10A_Gala
43,250,105

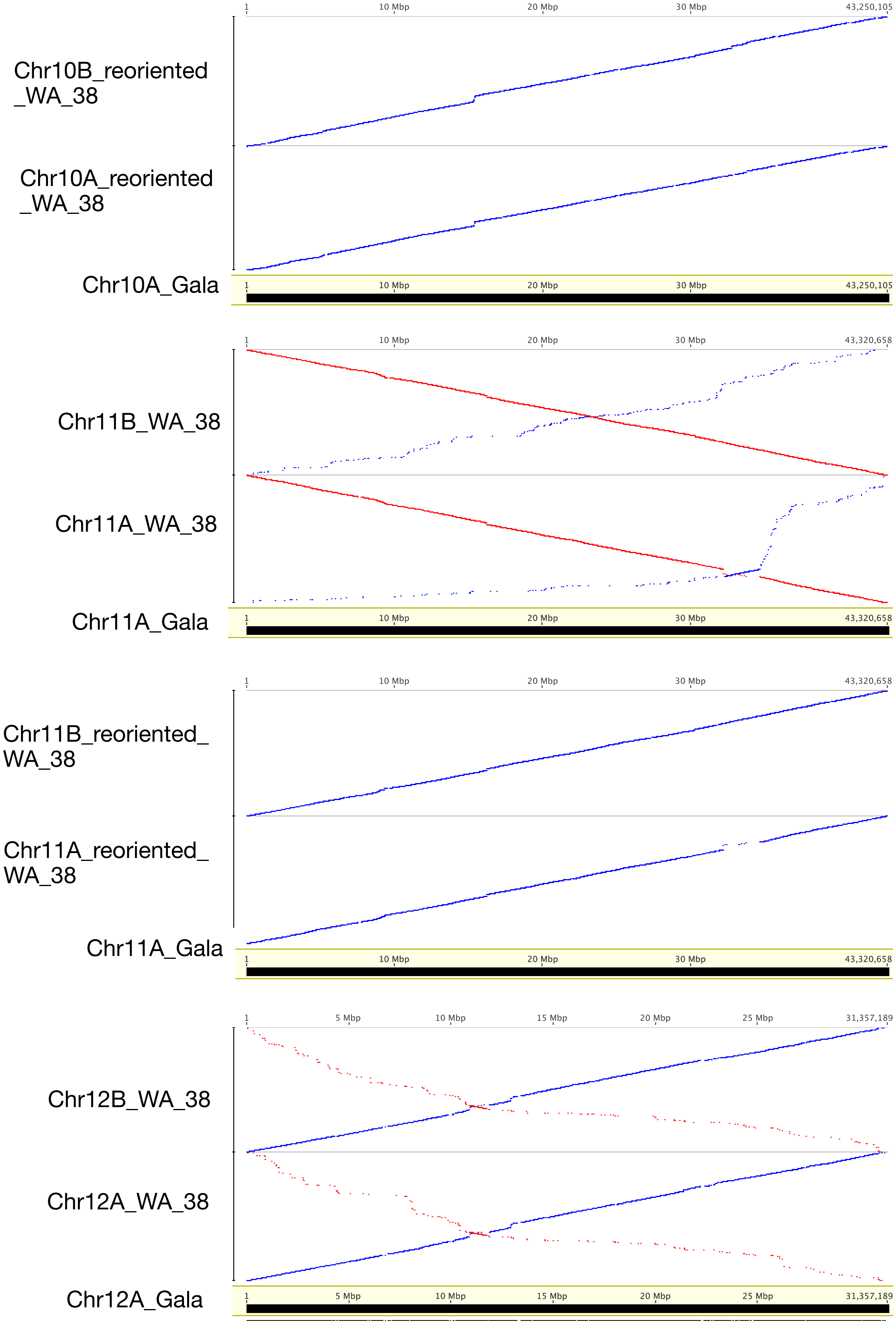
Chr10B_reoriented _WA_38
Chr10A_reoriented _WA_38
Chr10A_Gala
1
10 Mbp
20 Mbp
30 Mbp
43,250,105
Chr11B_WA_38
Chr11A_WA_38
Chr11A_Gala
43,320,658
Chr11B_reoriented_ WA_38
Chr11A_reoriented_ WA_38
Chr11A_Gala
Chr12B_WA_38
Chr12A_WA_38
Chr12A_Gala
5 Mbp
15 Mbp
25 Mbp
31,357,189

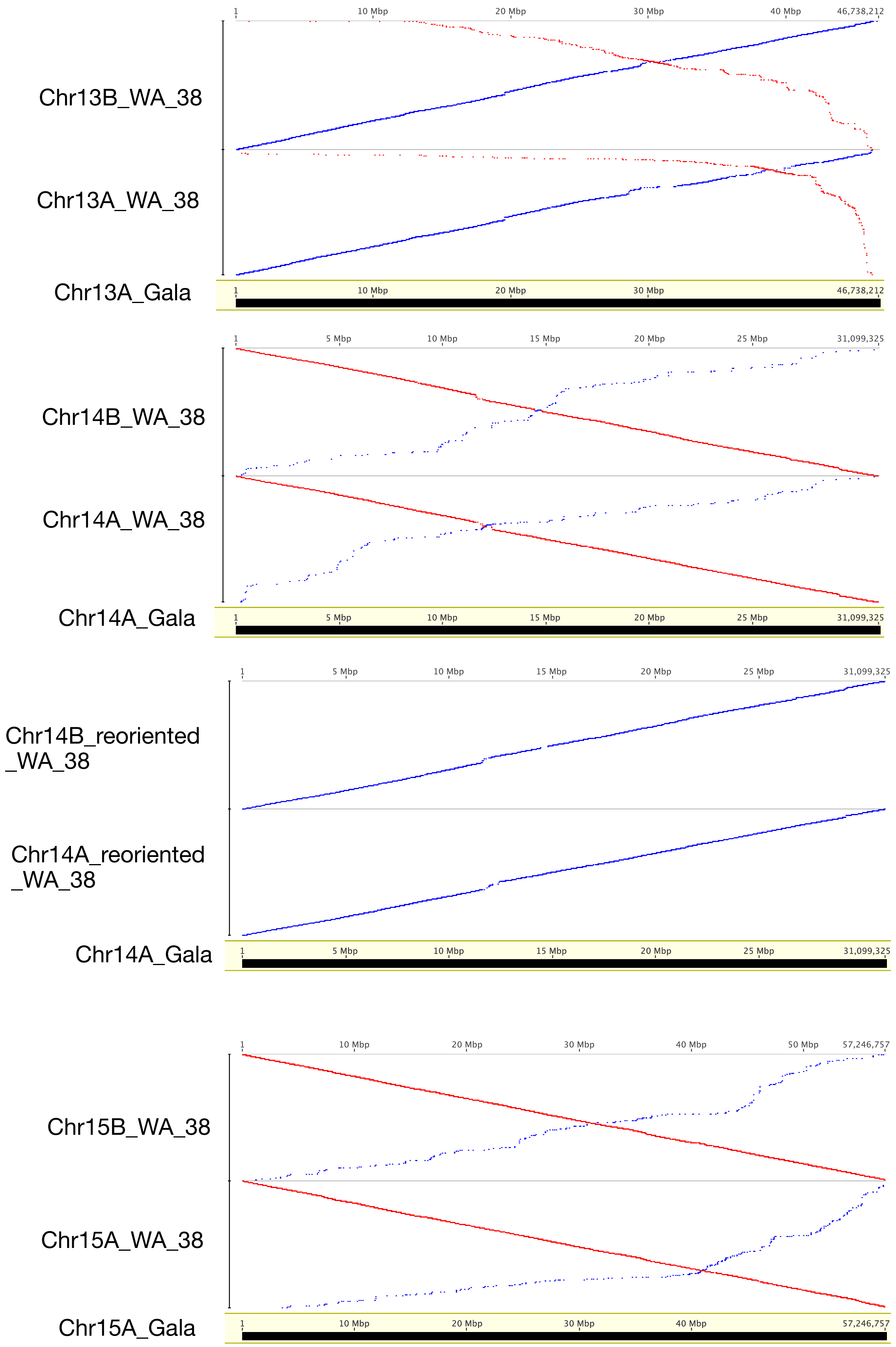

1
10 Mbp
20 Mbp
30 Mbp
40 Mbp
46,738,212
Chr13B_WA_38
Chr13A_WA_38
Chr13A_Gala
1
5 Mbp
10 Mbp
15 Mbp
20 Mbp
25 Mbp
31,099,325
Chr14B_WA_38
Chr14A_WA_38
Chr14A_Gala
Chr14B_reoriented _WA_38
Chr14A_reoriented _WA_38
Chr14A_Gala
1
10 Mbp
20 Mbp
30 Mbp
40 Mbp
50 Mbp
57,246,757
Chr15B_WA_38
Chr15A_WA_38
Chr15A_Gala

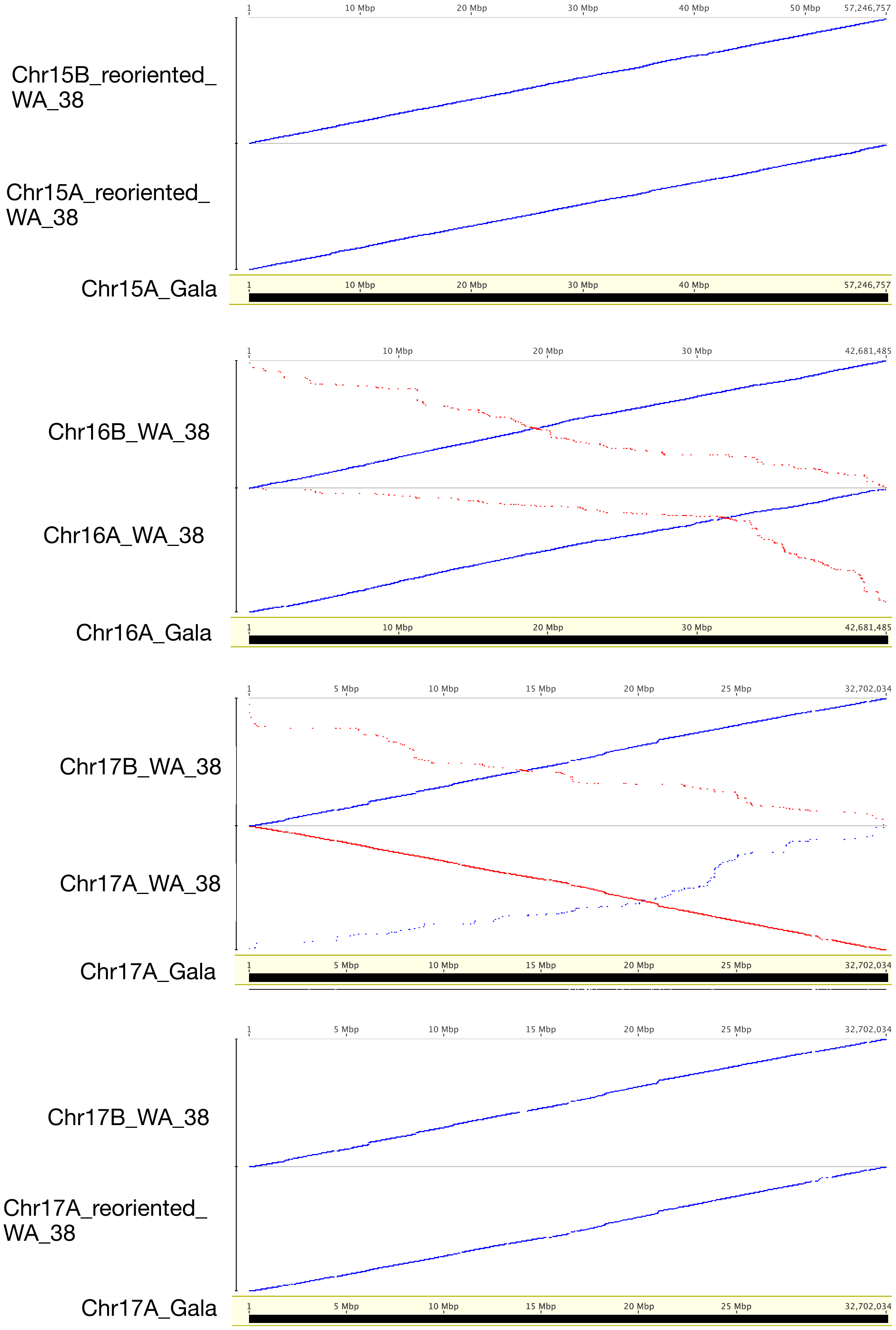

Chr15B_reoriented_WA_38
Chr15A_reoriented_WA_38
Chr15A_Gala
1
10 Mbp
20 Mbp
30 Mbp
40 Mbp
50 Mbp
57,246,757
Chr16B_WA_38
Chr16A_WA_38
Chr16A_Gala
42,681,485
Chr17B_WA_38
Chr17A_WA_38
Chr17A_Gala
5 Mbp
15 Mbp
25 Mbp
32,702,034
Chr17B_WA_38
Chr17A_reoriented_WA_38
Chr17A_Gala

Supplemental Figure 3. Hi-C contact map for Hap1 (**A**) and Hap2 (**B**) of 'WA 38' genome assembly.

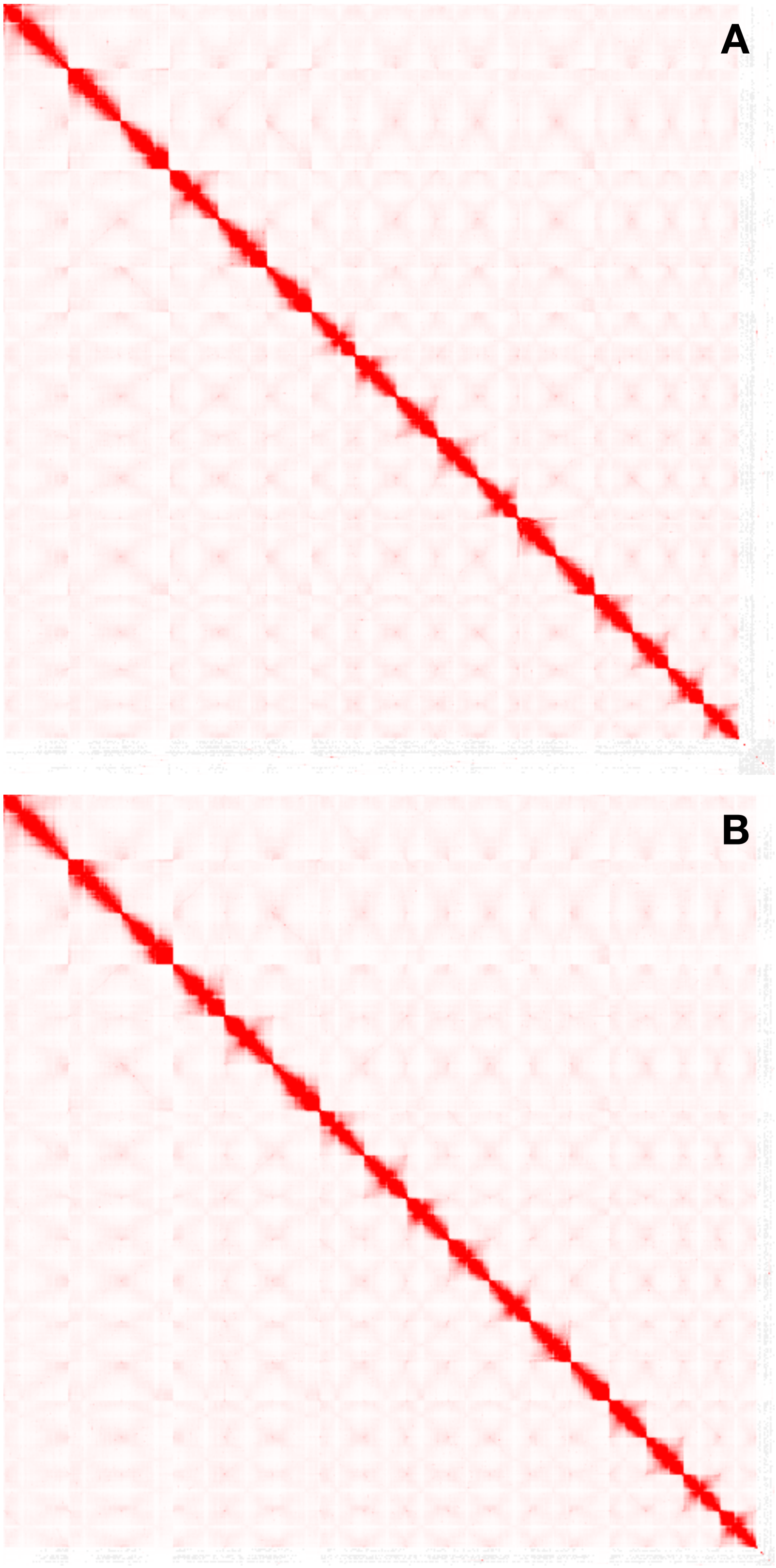

Supplemental Figure 4: Structural variation between reference (WA 38 hapA) and query (WA 38 hapB). (A), Size distribution of all variants of all sizes; (B), Size distribution of variants ranging from 50-500bp; (C), Size distribution of variants ranging from 500-10,000bp; (D), Cumulative sequence length of reference and query assemblies.

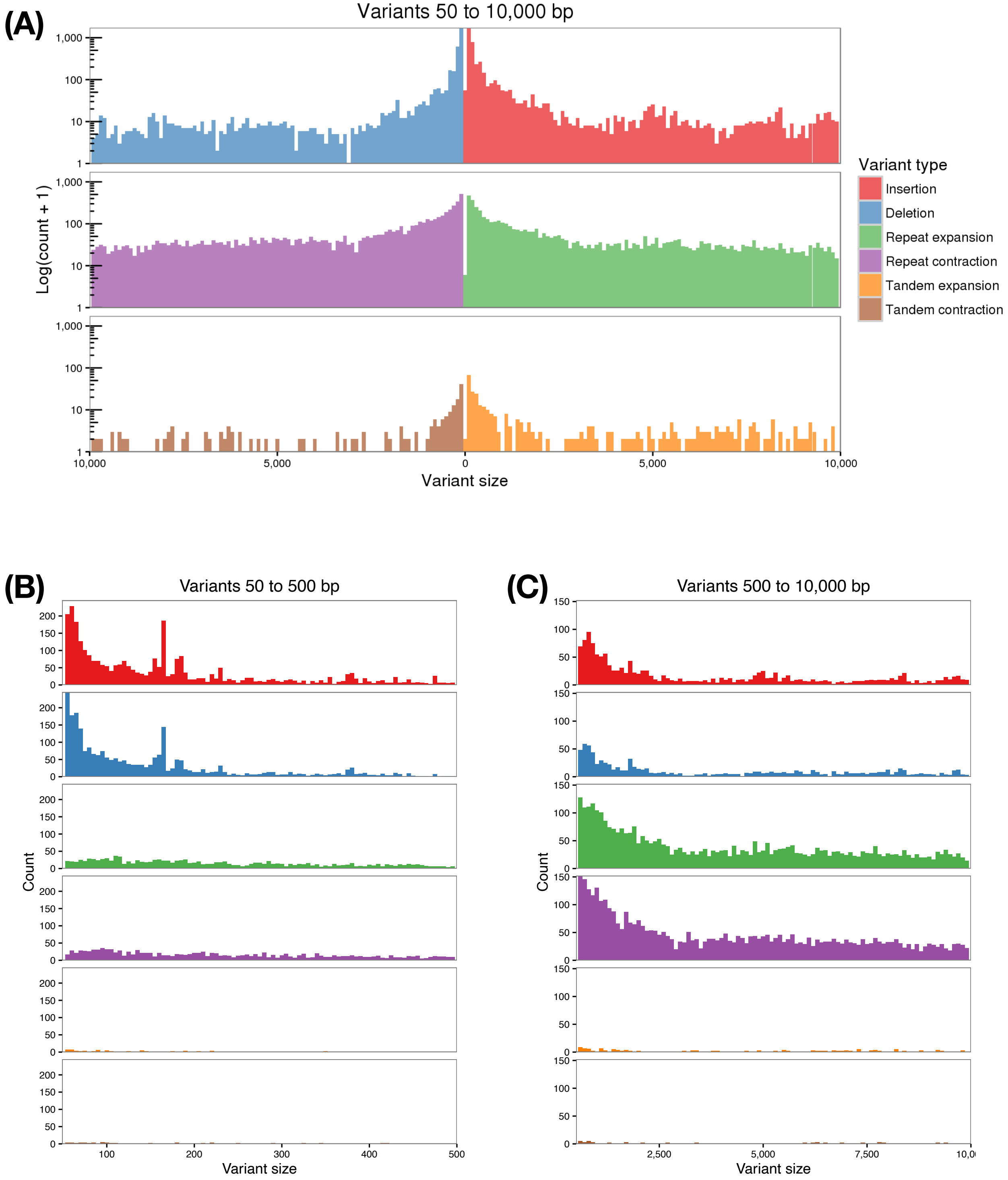

(D)

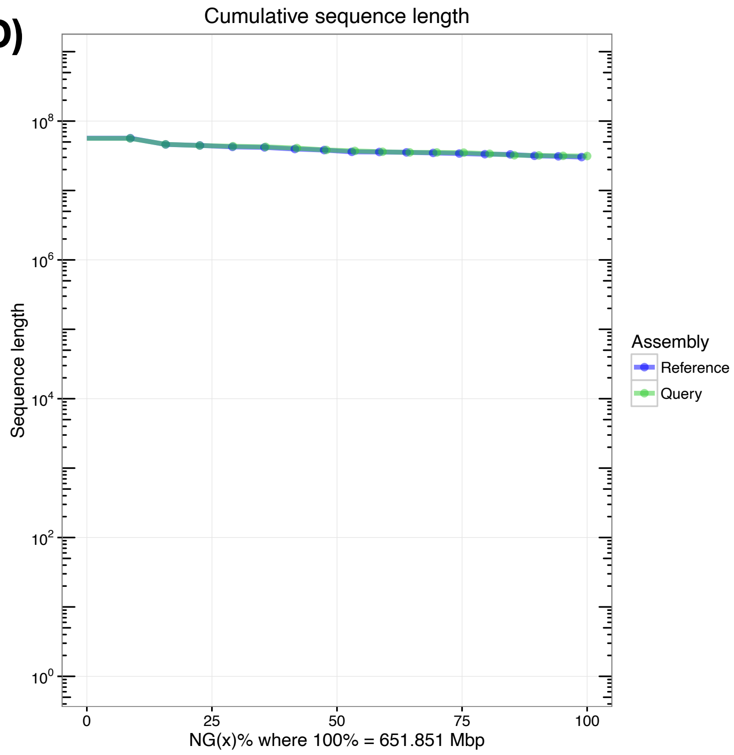
Cumulative sequence length
Sequence length
$10^8$
$10^6$
$10^4$
$10^2$
$10^0$
0
25
50
75
100
NG(x)% where 100% = 651.851 Mbp
Assembly
Reference
Query

Supplemental figure 5. Dot plots of genome alignment (**A**) and chromosome alignment (**B**) of the two ‘WA 38’ haplomes.

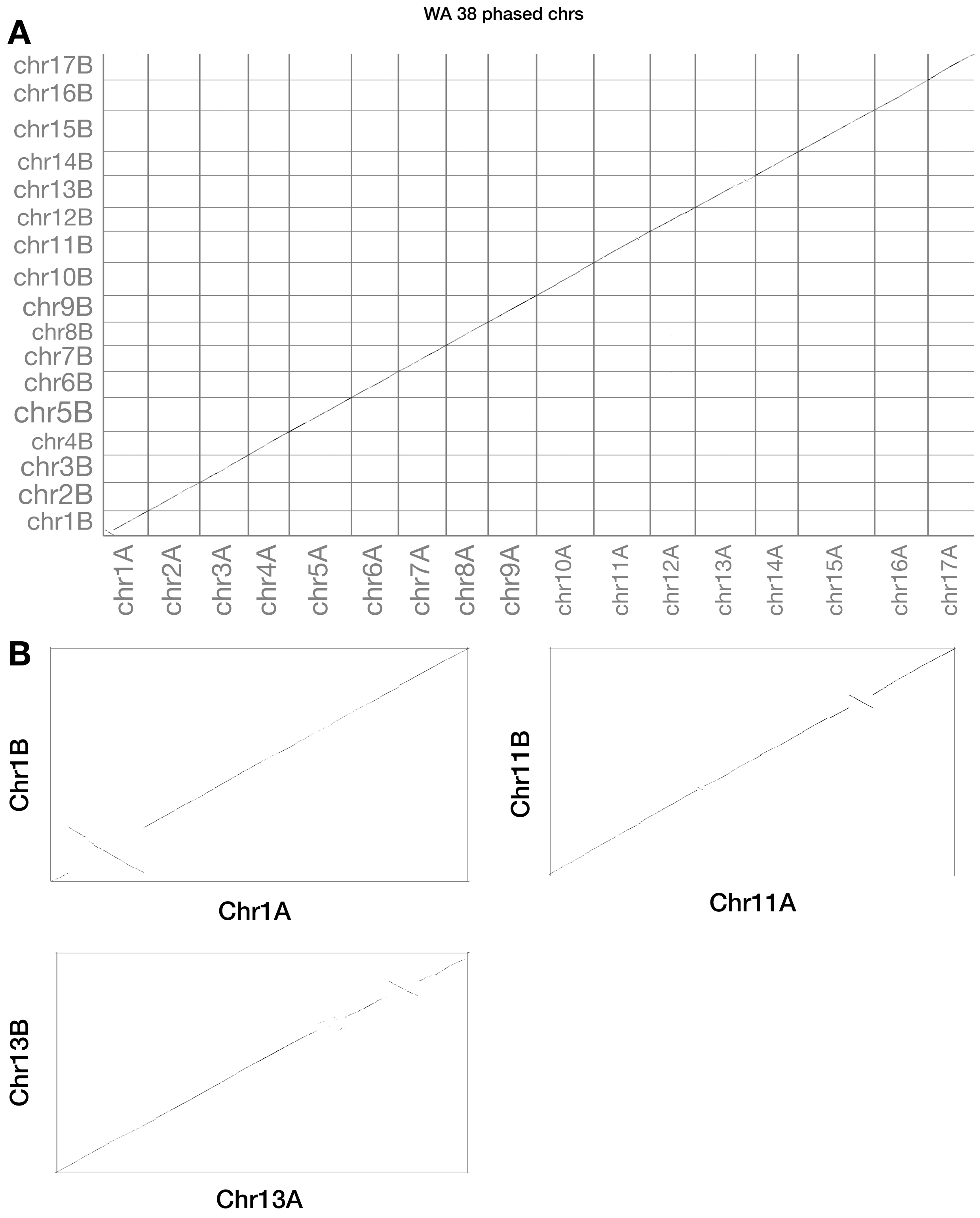

Supplemental Figure 6. Issues with BRAKER gene models. (A) shows an example of gene models overlaps with repeat region and splice variants of the same gene do not overlap. (B) shows an example of overlapping gene models on the same strand. Blue lines represent gene models and mRAN, orange lines represent repeat regions.

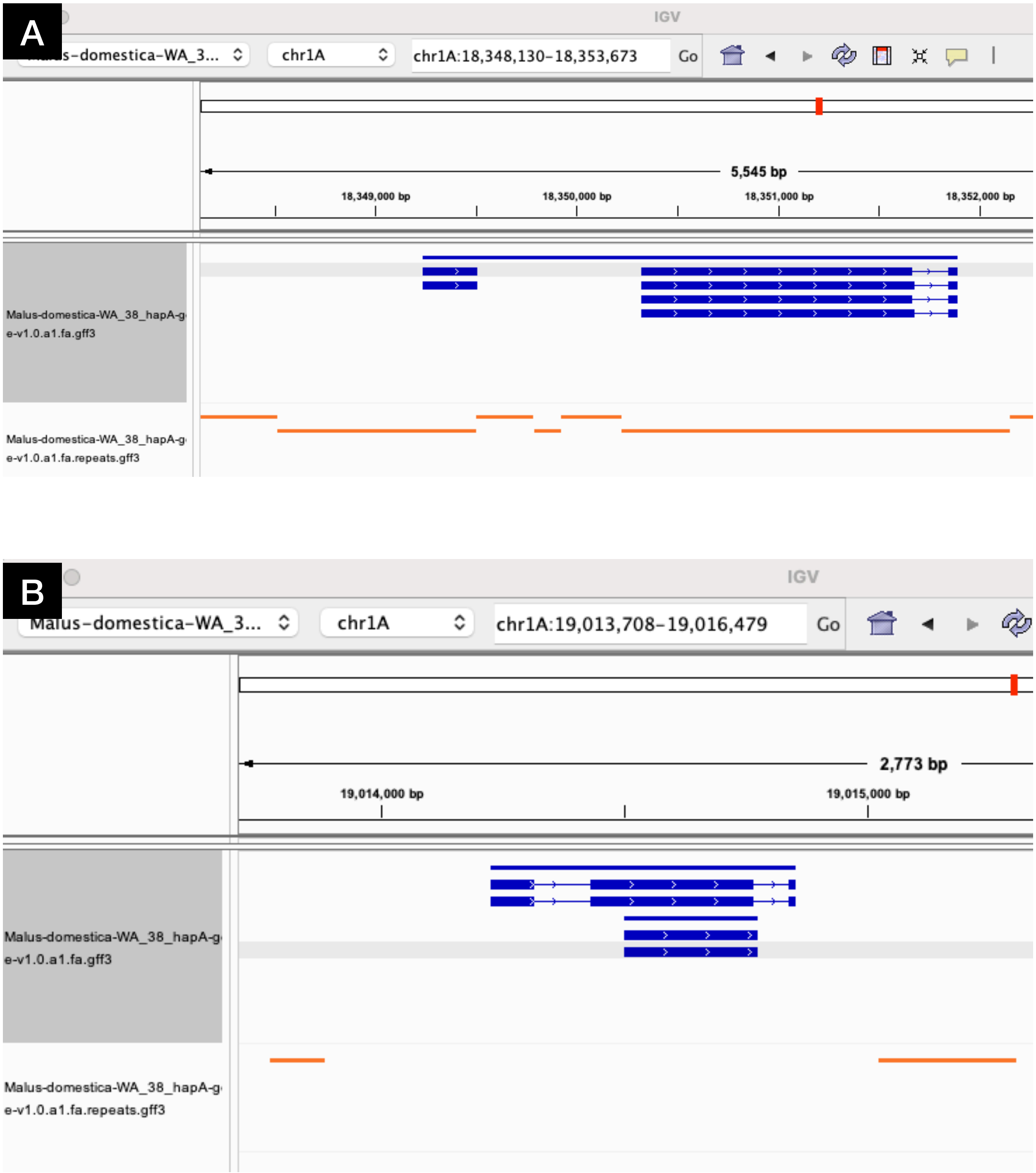

Supplemental Figure 7. Riparian plot comparing ‘WA 38’ Haplotype A and B with ‘Honeycrisp’ Haplotype A and B and ‘Golden Delicious’ (GDDH13) genomes by physical location.

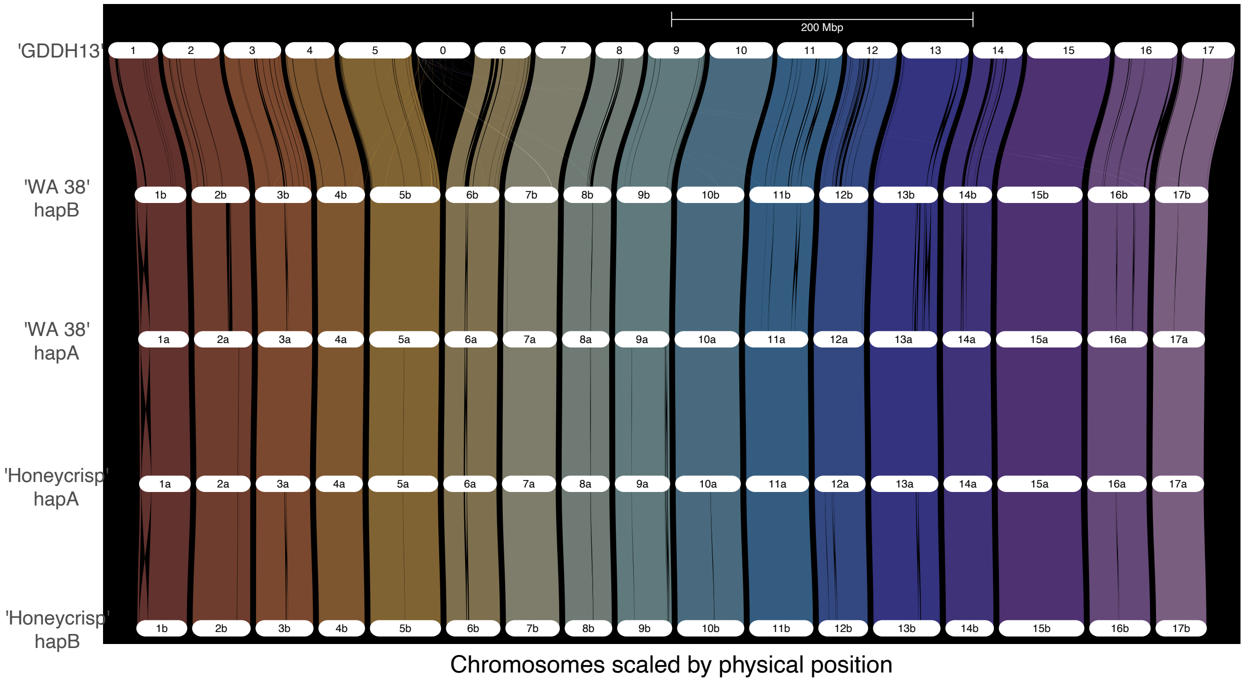

Supplemental Figure 8. CROG analysis using 'WA 38' Subset 1 (**A** & **C**) and Subset 3 (**B** & **D**). **A** & **B** are CROG gene count cluster maps. Each row represents a CROG and each column represents a genomes. Color indicates the number of genes in each cell relative to the row average (z-score). Warmer color indicates more genes. Cooler color indicates fewer genes. The darker a color, the closer the value is to the row average. Green boxes highlights shared 'cold' orthogroups among genomes released from the same publication, or analyzed by the same research groups/ individual. **C** & **D** are CROG gene count z-score box plots summarizing z-score distribution of CROG gene counts in selected pome fruit genomes. Genome and annotation abbreviations can be found in Supplemental Table 1.

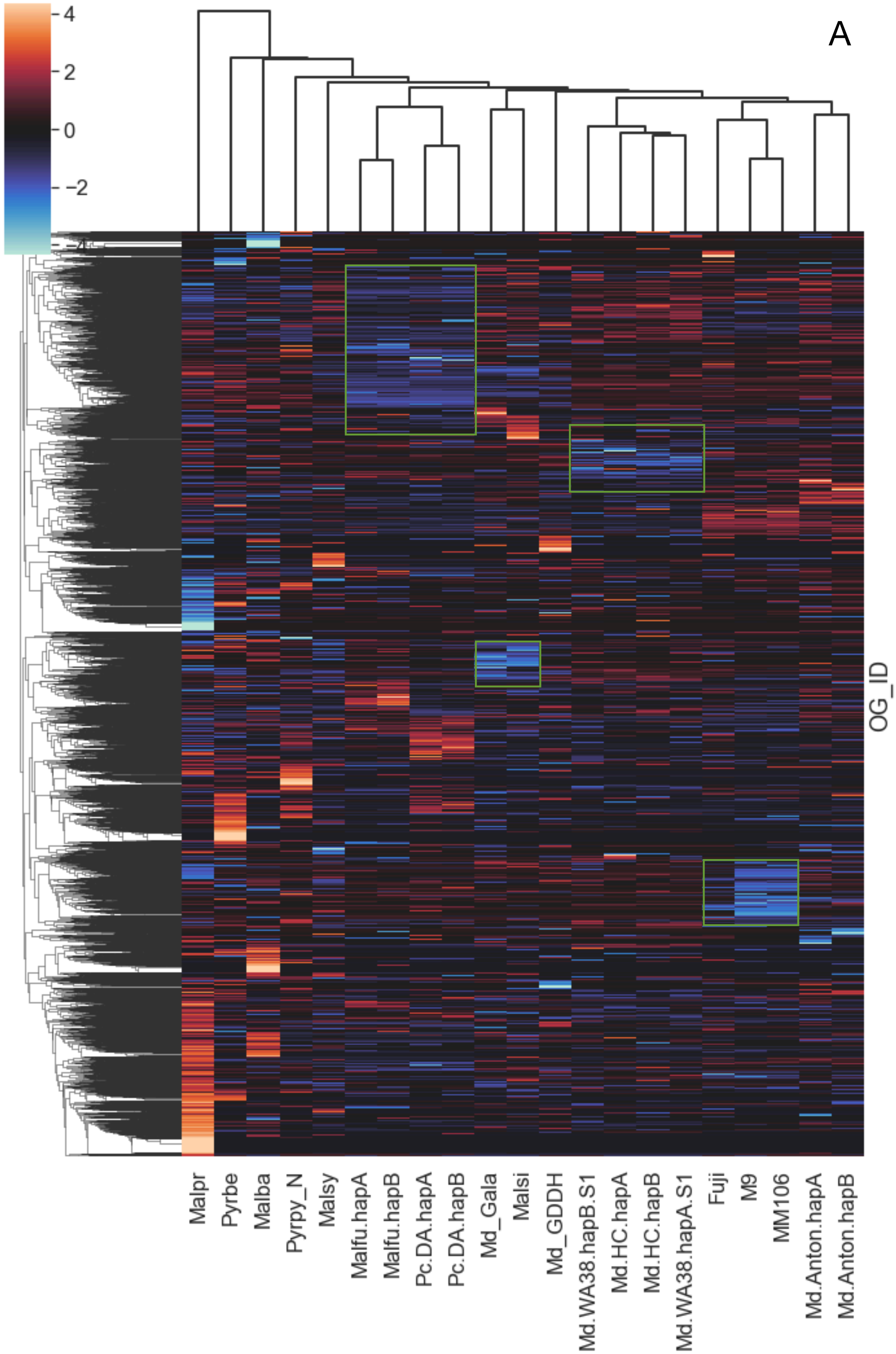

A
4
2
0
−2
−4
OG_ID
Malpr
Pyrbe
Malba
Pyrpy_N
Malsy
Malfu.hapA
Malfu.hapB
Pc.DA.hapA
Pc.DA.hapB
Md_Gala
Malsi
Md_GDDH
Md.WA38.hapB.S1
Md.HC.hapA
Md.HC.hapB
Md.WA38.hapA.S1
Fuji
M9
MM106
Md.Anton.hapA
Md.Anton.hapB

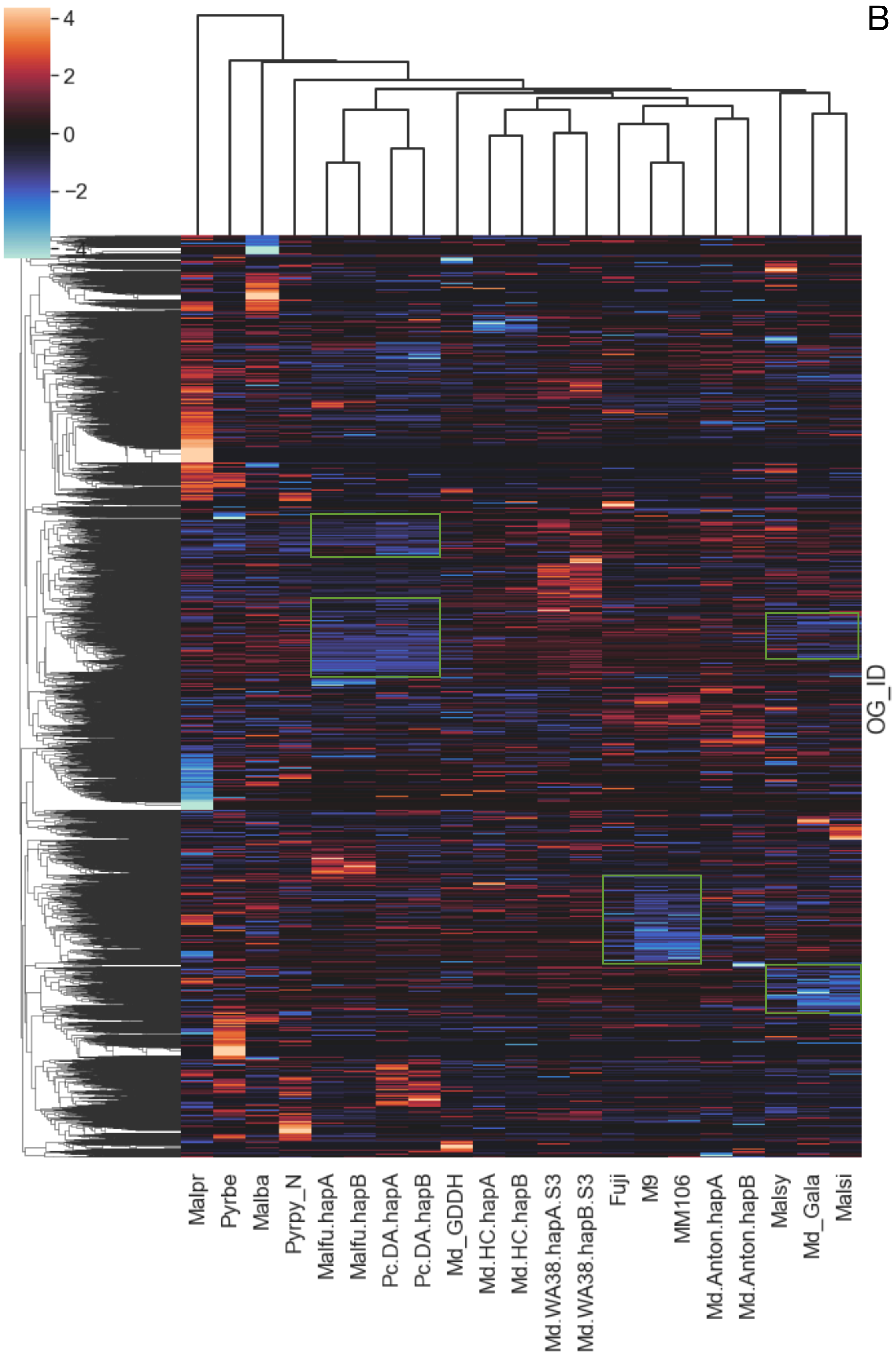

B
4
2
0
−2
−4
OG_ID
Malpr
Pyrbe
Malba
Pyrpy_N
Malfu.hapA
Malfu.hapB
Pc.DA.hapA
Pc.DA.hapB
Md_GDDH
Md.HC.hapA
Md.HC.hapB
Md.WA38.hapA.S3
Md.WA38.hapB.S3
Fuji
M9
MM106
Md.Anton.hapA
Md.Anton.hapB
Malsy
Md_Gala
Malsi

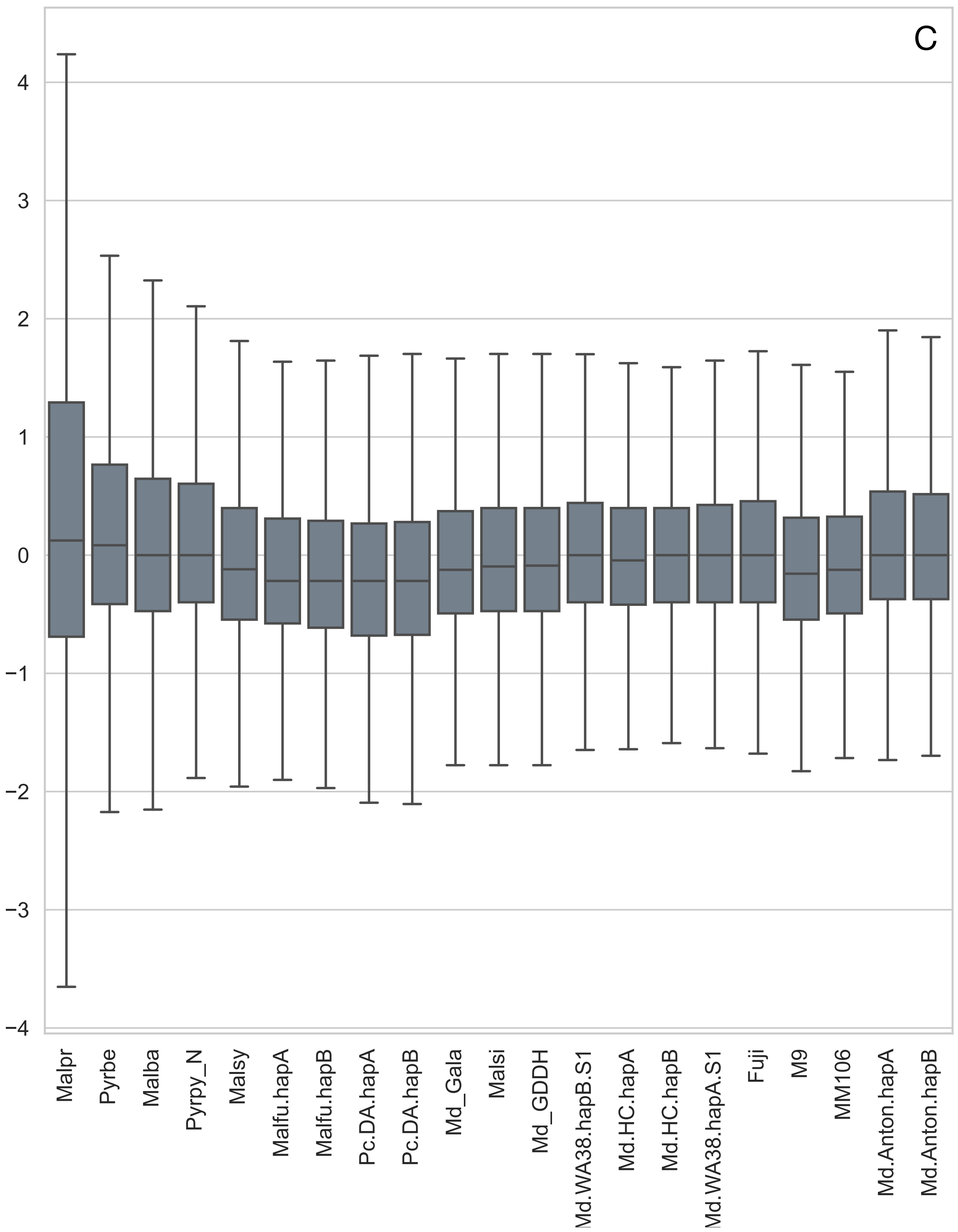

C
4
3
2
1
0
−1
−2
−3
−4
Malpr
Pyrbe
Malba
Pyrpy_N
Malsy
Malfu.hapA
Malfu.hapB
Pc.DA.hapA
Pc.DA.hapB
Md_Gala
Malsi
Md_GDDH
Md.WA38.hapB.S1
Md.HC.hapA
Md.HC.hapB
Md.WA38.hapA.S1
Fuji
M9
MM106
Md.Anton.hapA
Md.Anton.hapB

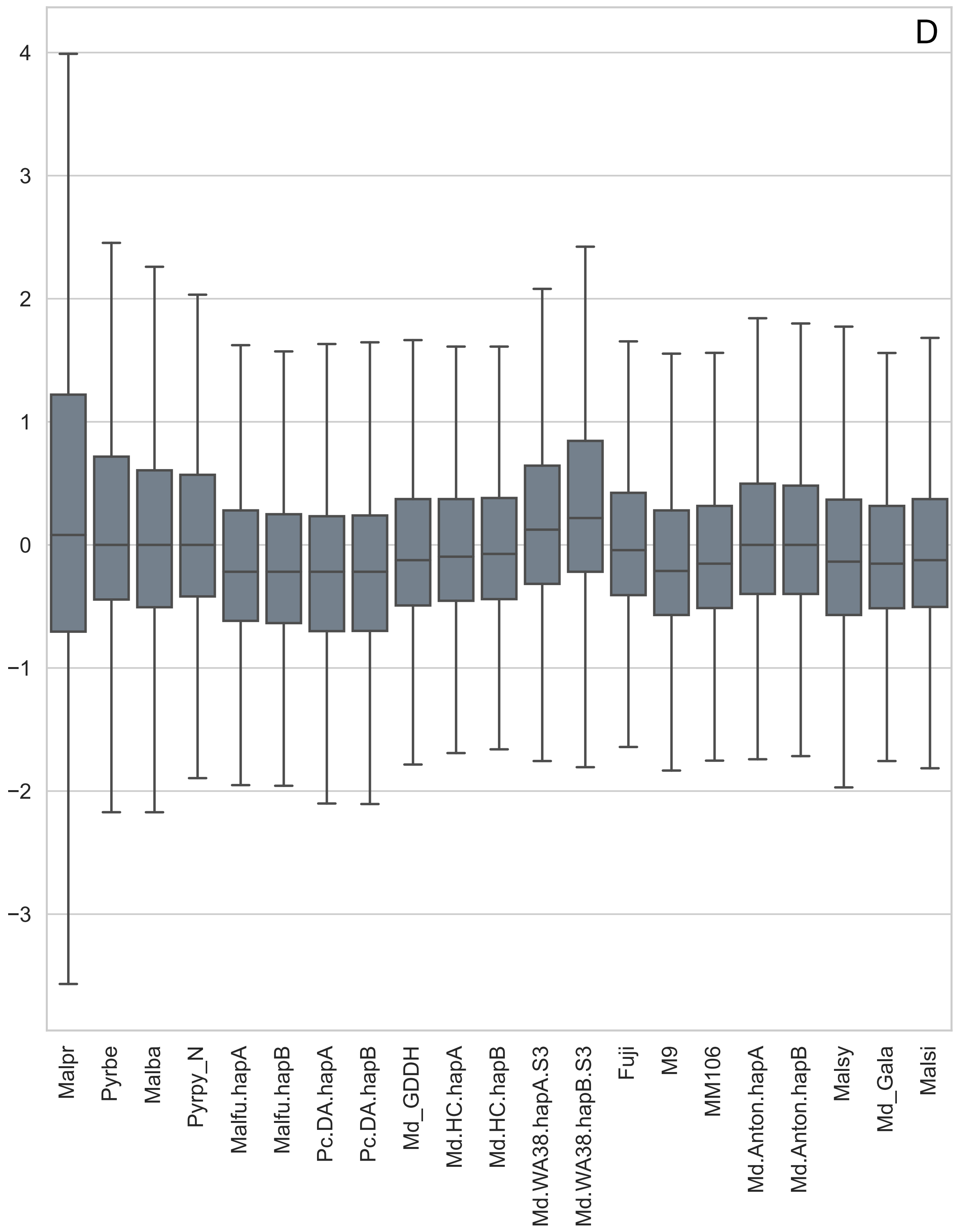

D
4
3
2
1
0
−1
−2
−3
Malpr
Pyrbe
Malba
Pyrpy_N
Malfu.hapA
Malfu.hapB
Pc.DA.hapA
Pc.DA.hapB
Md_GDDH
Md.HC.hapA
Md.HC.hapB
Md.WA38.hapA.S3
Md.WA38.hapB.S3
Fuji
M9
MM106
Md.Anton.hapA
Md.Anton.hapB
Malsy
Md_Gala
Malsi
